# Exploring archaeogenetic studies of dental calculus to shed light on past human migrations in Oceania

**DOI:** 10.1101/2023.10.18.563027

**Authors:** Irina M. Velsko, Zandra Fagernäs, Monica Tromp, Stuart Bedford, Hallie R. Buckley, Geoffrey Clark, John Dudgeon, James Flexner, Anatauarii Leal-Tamarii, Cecil M. Lewis, Elizabeth Matisoo-Smith, Kathrin Nägele, Andrew T. Ozga, Adam B. Rohrlach, Cosimo Posth, Richard Shing, Matthew Spriggs, Edson Willie, Frédérique Valentin, Christina Warinner

**Affiliations:** Department of Archaeogenetics, Max Planck Institute for Evolutionary Anthropology, Leipzig, Germany; Department of Archaeology, Max Planck Institute for the Science of Human History, Jena, Germany; Southern Pacific Archaeological Research, Archaeology Programme, University of Otago, Dunedin, New Zealand; Department of Anatomy, School of Biomedical Sciences, University of Otago, Dunedin, New Zealand; Department of Archaeology and Natural History, College of Asia and the Pacific, The Australian National University, Canberra, Australia; Department of Linguistic and Cultural Evolution, Max Planck Institute for Evolutionary Anthropology, Leipzig, Germany; Department of Anthropology, Idaho State University, Pocatello, ID, USA; Archaeology, School of Humanities, University of Sydney, Sydney, Australia; Direction de la Culture et du Patrimoine, Puna’auia, French Polynesia; Department of Anthropology, University of Oklahoma, Norman, OK, USA; Department of Biological Sciences, Halmos College of Arts and Sciences, Nova Southeastern University, Fort Lauderdale, FL, USA; School of Computer and Mathematical Sciences, The University of Adelaide, Adelaide, Australia; Archaeo- and Palaeogenetics, Institute for Archaeological Sciences, Department of Geosciences, University of Tübingen, Tübingen 72074, Germany; Senckenberg Centre for Human Evolution and Palaeoenvironment at the University of Tübingen, Tübingen 72074, GermanyDepartment of Archaeo- and Palaeogenetics, Eberhard Karls Universität Tübingen, Tübingen, Germany; Vanuatu Cultural Centre, Port-Vila, Vanuatu; School of Archaeology and Anthropology, College of Arts & Social Sciences, The Australian National University, Canberra, Australia; CNRS UMR 8068, MSH Mondes, Nanterre, France; Faculty of Biological Sciences, Friedrich Schiller University, Jena, Germany; Department of Anthropology, Harvard University, Cambridge, MA, USA

**Keywords:** ancient dental calculus, oral microbiome, Oceania, migration, Pacific

## Abstract

The Pacific islands have experienced multiple waves of human migrations, providing a case study for exploring the potential of using the microbiome to study human migration. We performed a metagenomic study of archaeological dental calculus from 103 ancient individuals, originating from 12 Pacific islands and spanning a time range of ∼3000 years. Oral microbiome DNA preservation in calculus is far higher than that of human DNA in archaeological bone from the Pacific, and comparable to that seen in calculus from temperate regions. Variation in the microbial community composition was minimally driven by time period and geography within the Pacific, while comparison with samples from Europe, Africa, and Asia reveal the microbial communities of Pacific calculus samples to be distinctive. Phylogenies of individual bacterial species in Pacific calculus reflect geography. Archaeological dental calculus shows potential to yield information about past human migrations, complementing studies of the human genome.

## Introduction

Archaeogenetics studies of ancient human migrations are conventionally conducted by analyzing human DNA from skeletal elements, such as teeth and the petrous portion of the temporal bone. Human population movements across the Pacific have been reconstructed in this manner, revealing successive waves of migration during the Pleistocene and Holocene^1–5^. However, ancient DNA work in the Pacific is challenging as the studied populations are closely related and migrations take place over short periods of time. Further, the high temperatures and humidity generally increase the rate of DNA decay^6,7^, making human DNA preservation highly variable^1^. Ancient dental calculus, the calcified oral biofilm preserved on teeth, offers a possible alternative approach to studying human migrations in the Pacific, which has not yet been widely explored^8,9^.

Dental calculus forms when the bacterial biofilm known as dental plaque naturally calcifies on the surfaces of teeth. The dental plaque microbes become encased within the mineral matrix, preserving biomolecules including DNA, proteins, and small molecule metabolites, and this enables studies of oral bacterial communities stretching thousands of years back in time^10^. Because microbes have shorter generation times than humans, they offer the possibility to study the migrations of closely related populations over shorter timescales. Further, DNA within dental calculus is generally better preserved than DNA in skeletal tissues from the same individual^11^, making it a promising substrate to study in areas where skeletal DNA preservation is generally poor, such as the tropics.

The prospect of studying human migration through archaeological dental calculus presents several potential advantages, as it would allow for a holistic study of an individual’s life through just one sample - from migration and diet, to health and disease, and even occupational activities^12^. Dental calculus may prove to be an especially valuable study material in the Pacific, given the speed of settlement of the region and significant cultural changes over a short time. However, no study has to date attempted to investigate past human migrations through the oral microbiome, although the prospect has been discussed^8,9^.

Here, we explore the possibility of using archaeological dental calculus to study past human migrations, using the Pacific islands as a case study. Shotgun metagenomic sequencing was performed on a total of 103 dental calculus samples from 12 Pacific islands, spanning a time range of nearly 3000 years (Table 1). We show that preservation is variable, but that most samples have a well-preserved oral microbiome, providing support to the exceptional preservation of dental calculus, even in challenging environments. We find that variation in dental calculus microbial community composition does not have a clear geographic or temporal structure, but rather may be influenced by local factors specific to each island. In contrast, phylogenetic analyses of individual oral bacterial taxa exhibit temporal trends, but the ability to detect this signal depends on the selected species’ prevalence and abundance. Overall, we find that metagenomic analysis of archaeological dental calculus has the potential to reveal information about past human migration patterns. When combined with human DNA analysis and other approaches, such as paleodietary studies using palaeoproteomics and microremains, it promises to enrich our understanding of the dynamic biological and cultural processes that accompanied past migrations in the Pacific islands.

**Table 1.**
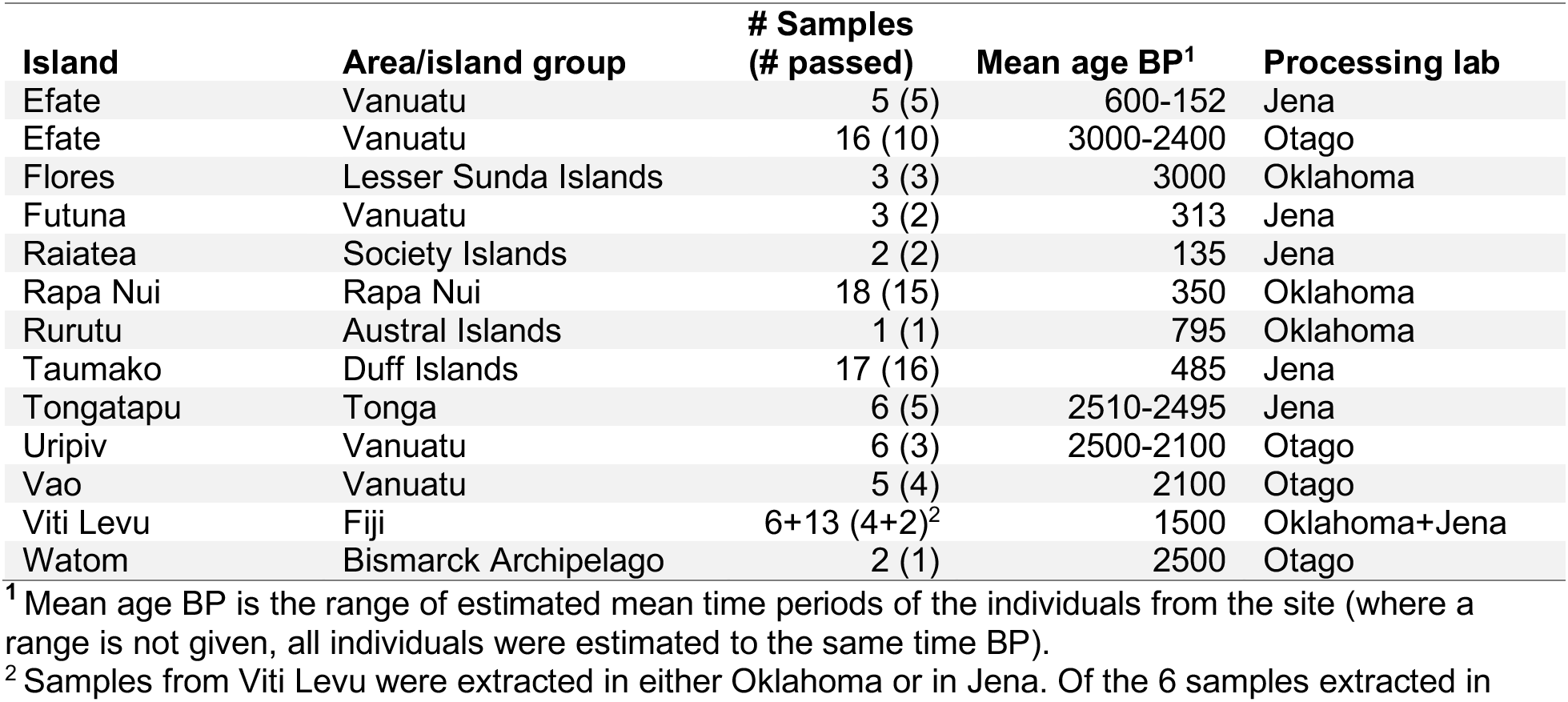
Archaeological dental calculus samples included in this study.

## Results

### Preservation of endogenous ancient DNA

Obtaining well-preserved ancient DNA is a persistent challenge in archaeogenetic studies of the tropics, as both temperature and humidity contribute to DNA decay^6,7^. Using SourceTracker analysis^13^, we estimated the proportion of microbial taxa originating from endogenous and contaminant sources for each dental calculus sample in this study (Figure 1A). For 73 of the 103 archaeological dental calculus samples, at least 50% of microbial content is estimated to originate from an oral microbiome source, indicating good preservation of the dental calculus for these samples, with minimal contamination from exogenous sources. We further assessed the proportion of endogenous oral species using cuperdec^10^ (Supplemental figure S1) and found high consistency between estimated sample preservation using both methods. A PCA of well-preserved samples shows minimal overlap between dental calculus and the source samples used in SourceTracker, with calculus samples clustering distinctly from all sources (Figure 1B). All samples that passed the cuperdec threshold for preservation, 73 samples, were carried forward for analysis (Figure 1C).

**Figure 1.**
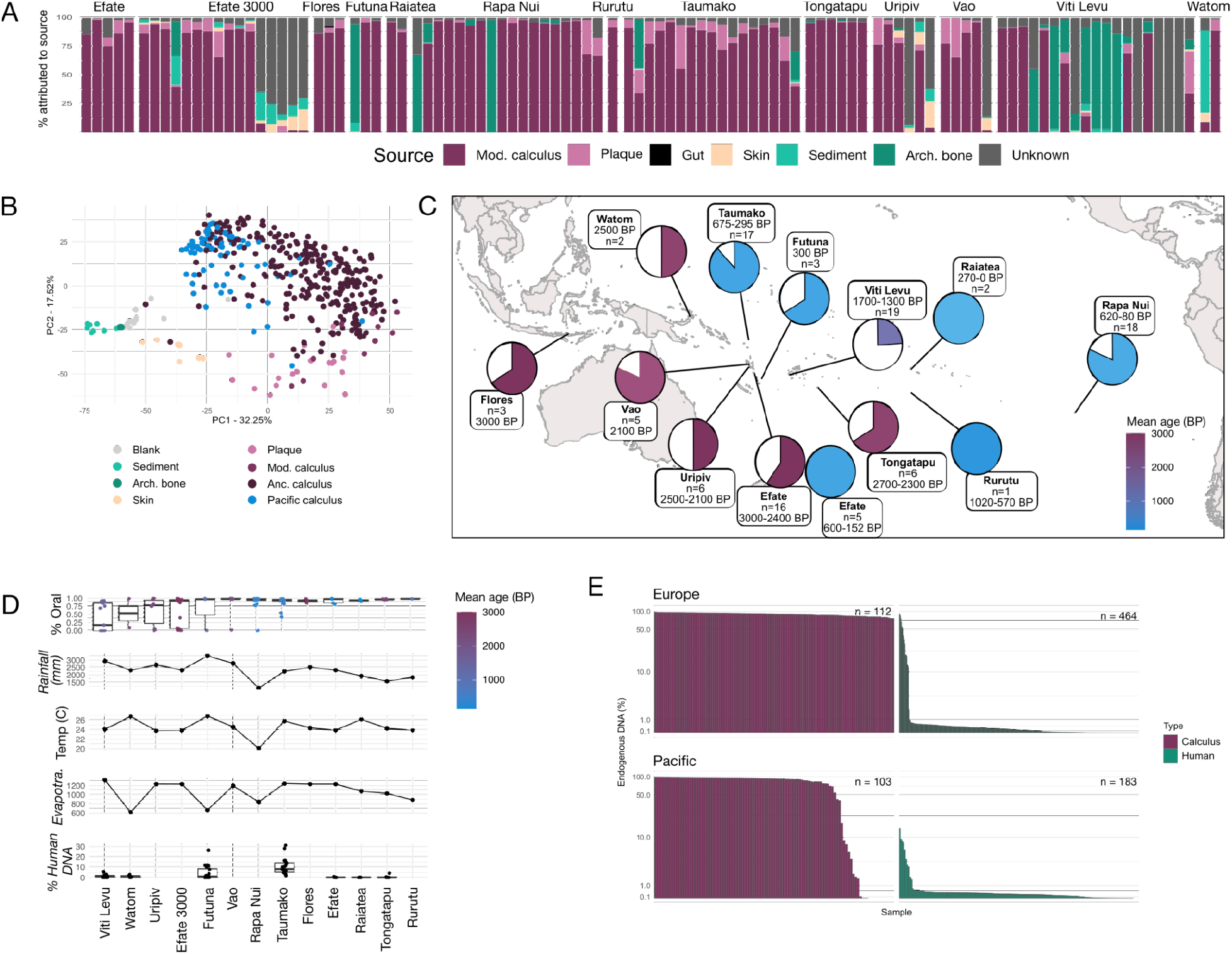
Preservation assessment of dental calculus samples. (**A**) SourceTracker analysis of species tables. Each bar represents a sample, colored by the proportion of each contributing source (sources are the same as in (B). (**B**) PCA of well-preserved calculus from the Pacific islands (blue) with samples from the same sources used in SourceTracker analysis plus additional ancient calculus. (**C**) Map of the islands from which samples were collected for this study, with the island name, number of collected samples, and age of the site. Pie charts indicate the fraction of samples from each site that were considered well-preserved, and are colored by age of the site. (**D**). Comparison of endogenous oral bacterial DNA and human host DNA in samples with environmental conditions on the islands that may affect DNA preservation. No associations between preservation of either oral bacterial DNA or human host DNA were found with any of the measured environmental factors. (**E**). Comparison between the Pacific islands and northern Europe (England/the Netherlands) of the percentage of endogenous oral microbiome DNA in ancient dental calculus and human DNA in bones/teeth. The number of samples in each group is indicated next to the bars. Ancient dental calculus contains high levels of endogenous oral bacterial DNA in the Pacific islands similar to that seen in northern Europe, in contrast to the lower levels of preserved human DNA in bones/teeth from the Pacific islands compared to northern Europe. Arch. bone - archaeological bone; Anc. calculus - ancient calculus; Mod. calculus - modern calculus.

The effect of sample age and environmental variables (average rainfall, average temperature, and average evapotranspiration of each island) on the proportion of taxa that could be assigned to oral sources (dental plaque and dental calculus), as well as the proportion of human DNA recovered from calculus, was investigated using beta regression (Figure 1D). Only sample age was a significant predictor of preservation (p=0.046, R^2^=0.083), yet the low R^2^ value indicates that additional, unknown factors more strongly affect preservation. Local environmental factors at the burial site and for each individual grave, such as soil type, humidity, and pH, are plausible candidates, but such data are not available.

Lastly, because the environment of the Pacific is less conducive to DNA preservation than colder, drier climates, we compared preservation patterns between the Pacific and northern Europe, and we asked whether the relative preservation of endogenous oral microbiome DNA in dental calculus is greater than that of human DNA in archaeological bone/teeth (Figure 1E). For the analysis, we considered all oral microbial DNA identified in dental calculus to be endogenous, and all human DNA present in bone/teeth to be endogenous. Overall, we find that endogenous preservation of ancient dental calculus in the Pacific is high, in which 72/103 samples (70%) are estimated to derive 80% or more of their composition from an oral microbiome source. This is only somewhat lower than that estimated for dental calculus from northern Europe, where 106/112 samples (95%) are estimated to derive more than 80% of their composition from an oral microbiome source. In contrast, human DNA preservation in skeletal material was much lower for both the Pacific and northern Europe, with 5/183 (2.7%) and 19/464 (4.1%) samples having endogenous DNA content of at least 5%, respectively, for samples in which ancient human DNA was detected. These results suggest that DNA within calculus is less prone to degradation and exogenous contamination than DNA in archaeological skeletal remains, and may be a more promising avenue of study for sites in warmer, more humid climates.

### Microbial community composition

We next asked whether the microbial community of well-preserved Pacific calculus samples showed temporal or spatial differences. If present, such patterns would suggest that the calculus microbiome changed with human migration through the Pacific. We performed a beta-diversity analysis and visualized the samples using PCA (Figure 2A). Testing the goodness of fit of sample cluster numbers to the data indicated that a single cluster optimally described the data (Supplemental Figure S2). The samples did not tightly cluster based on island or time period, nor did they plot along a cline that might suggest temporal or geographic change. PERMANOVA determined that the processing lab (R^2^ = 0.0275, F = 2.748, p = 0.01), island (R^2^ = 0.1913, F = 1.736, p = 0.001), and average GC content (R^2^ = 0.0508, F = 5.068, p = 0.002) were the most influential factors in how the samples plotted. When controlling for the lab in which samples were processed, island (R^2^ = 0.2045, F = 1.650, p = 0.035) and average GC content (R^2^ = 0.0761, F = 7.372, p = 0.01) were still significant drivers of the sample community composition. The average GC content of calculus is known to increase with sample age ^11,14,15^ through the taphonomic loss of AT-rich DNA fragments, and most islands are represented by samples from a single time period, therefore tying island and age/GC content.

**Figure 2.**
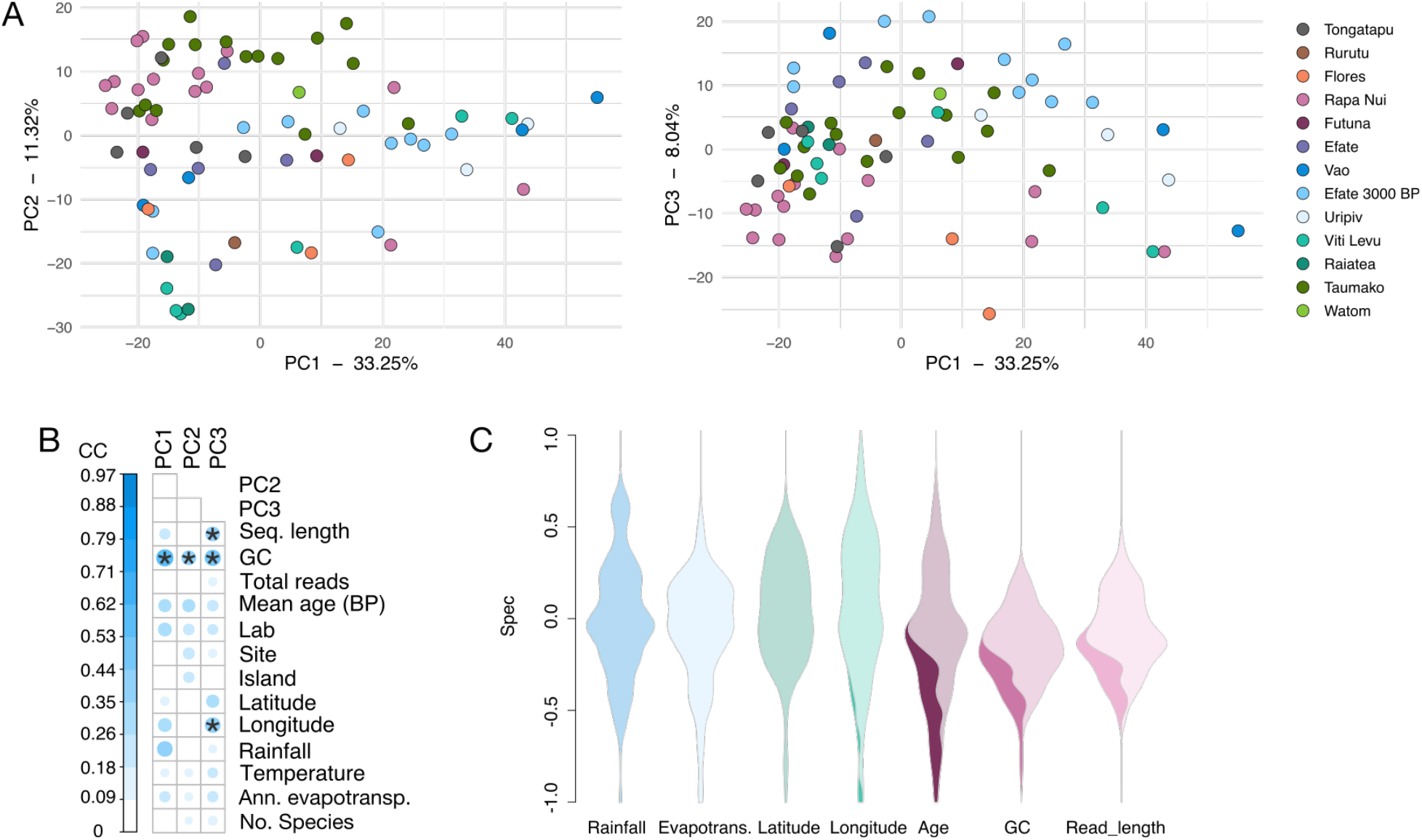
Community species profiles are minimally structured by island. (**A**) PCA of well-preserved calculus samples, colored by the island from which samples were collected; PCs 1, 2, and 3 are shown, accounting for >50% of the variation in the dataset. (**B**) Canonical correlation (CC) analysis comparing the positions of calculus samples in the PCA shown in (**A**) to environmental and laboratory metadata. Pearson correlation tests were used to determine if the correlations were significant. Metadata with a p ≤ 0.01 and CC value ≥ 0.4 are marked with an asterisk (*). Ann. evapotransp. - annual evapotranspiration; No. species - number of species. (**C**) Specificity of species in calculus samples to island conditions (blue), island location (green), or sequenced library characteristics (red). Dark colors in the violin plots indicate the proportion of species that are significantly associated with that metadata type. No dark color indicates that no species were specifically associated with that metadata.

However, the extent to which loss of AT-rich fragments affects species profiles, perhaps skewing older samples to have higher proportions of high-GC taxa, has not yet been extensively explored. We found that the species with the strongest PC1 loadings (Supplemental Table S3) suggested a taxonomic gradient relating to oxygen tolerance, such as that previously described in calculus samples from the archaeological site of Middenbeemster in the Netherlands^15^, but there was no clear association with GC content.

Low taxonomic assignment rates for the Pacific samples and the abundance of non-typical oral taxa that predominantly drive separation along PC1 suggest that taxonomic diversity in these samples may not be represented in the genomic databases we used for classification. We performed further taxonomic profiling with additional tools and databases, but were unable to resolve these issues, highlighting the difficulty of disentangling undescribed diversity from intrinsic biases in ancient calculus datasets, such as very short average DNA read lengths (≤ 70bp) (Supplemental figures S3, S4).

We performed canonical correlation analysis between PC loadings, environmental metadata, and laboratory characteristics, to determine if preservation may be influencing the calculus community composition (Figure 2B). Sample loadings on PC1 and PC2 were significantly correlated with average GC content, but not with sample metadata. We additionally looked for particular species that were significantly associated with environmental and laboratory metadata using the R package specificity (Figure 2C). No species were significantly associated with rainfall, evapotranspiration, or latitude, while a few were associated with longitude. In contrast, numerous species were significantly associated with sample age, average GC content, and average read length, which are themselves correlated, with some species significantly associated with more than one of these conditions. These results suggest that the environmental conditions tested here have minimal influence on the reconstructed calculus microbiome community, and instead other unexplored factors may be more influential.

### Comparison with global ancient calculus microbiome profiles

As this is the first large ancient dental calculus dataset published from the Pacific, we next wanted to know whether the microbiome communities fall within the known variation of published dental calculus datasets from across the globe. We compared the species profiles of our Pacific calculus samples to those from Europe, Africa, and Asia using PCA (Figure 3A). The Pacific samples and those from Japan^16^ cluster at the same end of the plot, suggesting that this region may have a particular species profile characteristic. Sample clustering based on continent was significant by PERMANOVA (p < 0.01, F = 2.47, R2 = 0.03114), while tests of beta-dispersion between the continents found differences only between the Pacific and Asia (Figure 3B, p < 0.05). However, the large difference in sample numbers for each continent make these comparisons less reliable. The distribution of samples across PC1 in the PCA may be largely driven by the number of species in each sample, as there is a moderate correlation between PC1 value and species counts (Figure 3C). The lower average number of species detected in Pacific samples compared to other continents may be related to the lower average number of reads in each sample that could be assigned taxonomy (Supplemental Figure S5), further hinting at unexplored diversity in the Pacific calculus.

**Figure 3.**
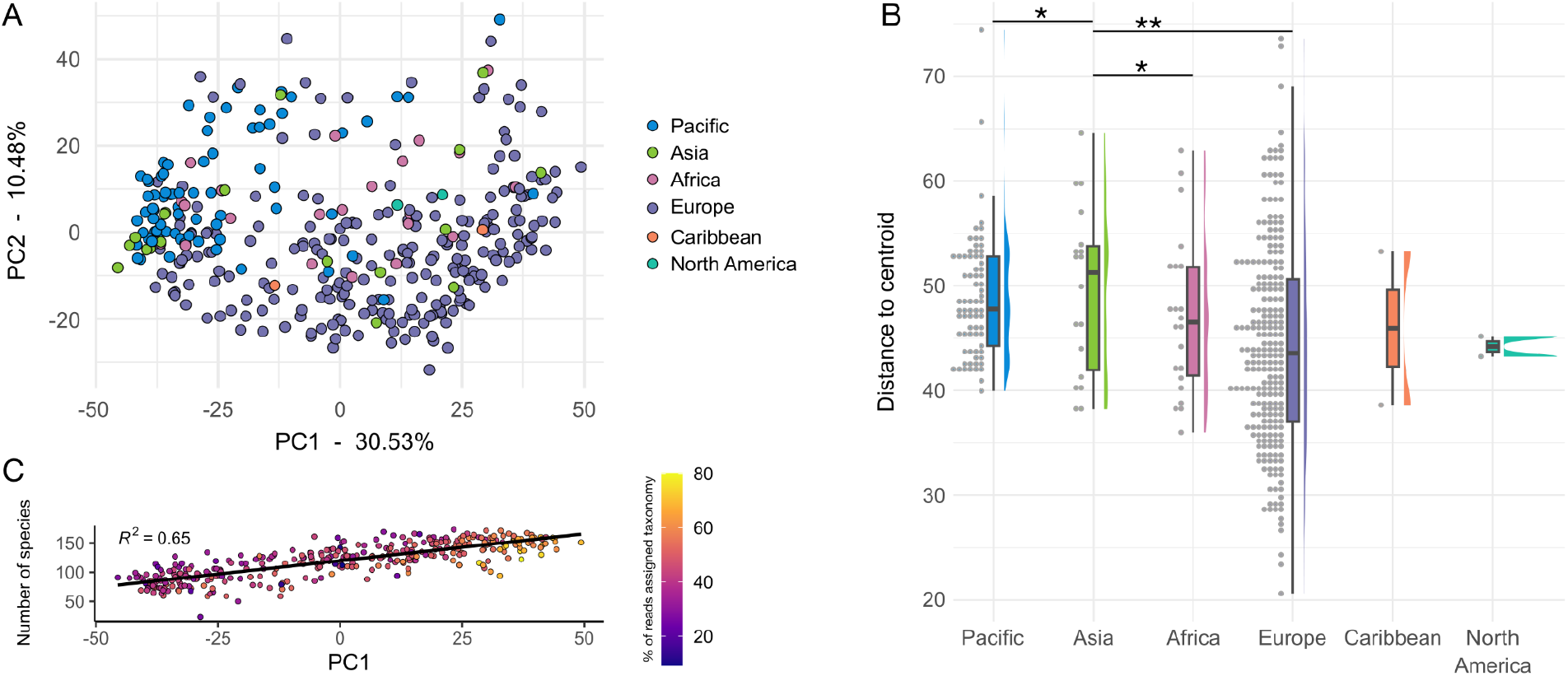
Situating the Pacific calculus samples within known ancient dental calculus microbial diversity. (**A**) PCA of Pacific calculus samples with ancient dental calculus from additional geographic regions. The additional samples are the same as those in Figure 1B. (**B**) The distance to the centroid of all samples in the PCA. * p < 0.05, ** p < 0.01. (**C**). The number of species in each sample, ordered by PC1 loading, and colored by the percentage of total reads in the sample that were assigned taxonomy. The trend line is fit with a generalized linear model.

### Gene content

We next investigated whether microbial gene content distinguishes the Pacific calculus samples from those of other continents. Hierarchical clustering of the KEGG orthologs (KOs) detected in samples did not cluster samples by continent or sample age (Figure 4A), the processing lab, or the study in which they were first presented (Supplemental Figure S6). The samples form two clusters (Clusters A and B) that correspond to their placement along PC1 in a PCA based on KO abundance (Supplemental Figure S7A) and based on species abundance (Supplemental Figure S7B), which is loosely correlated with the number of detectable KOs and species in each sample (Supplemental Figure S7C). Most of the Pacific samples fall in Sample Cluster A, which on average has lower species and KO counts (Supplemental Figure S7D, E).

**Figure 4.**
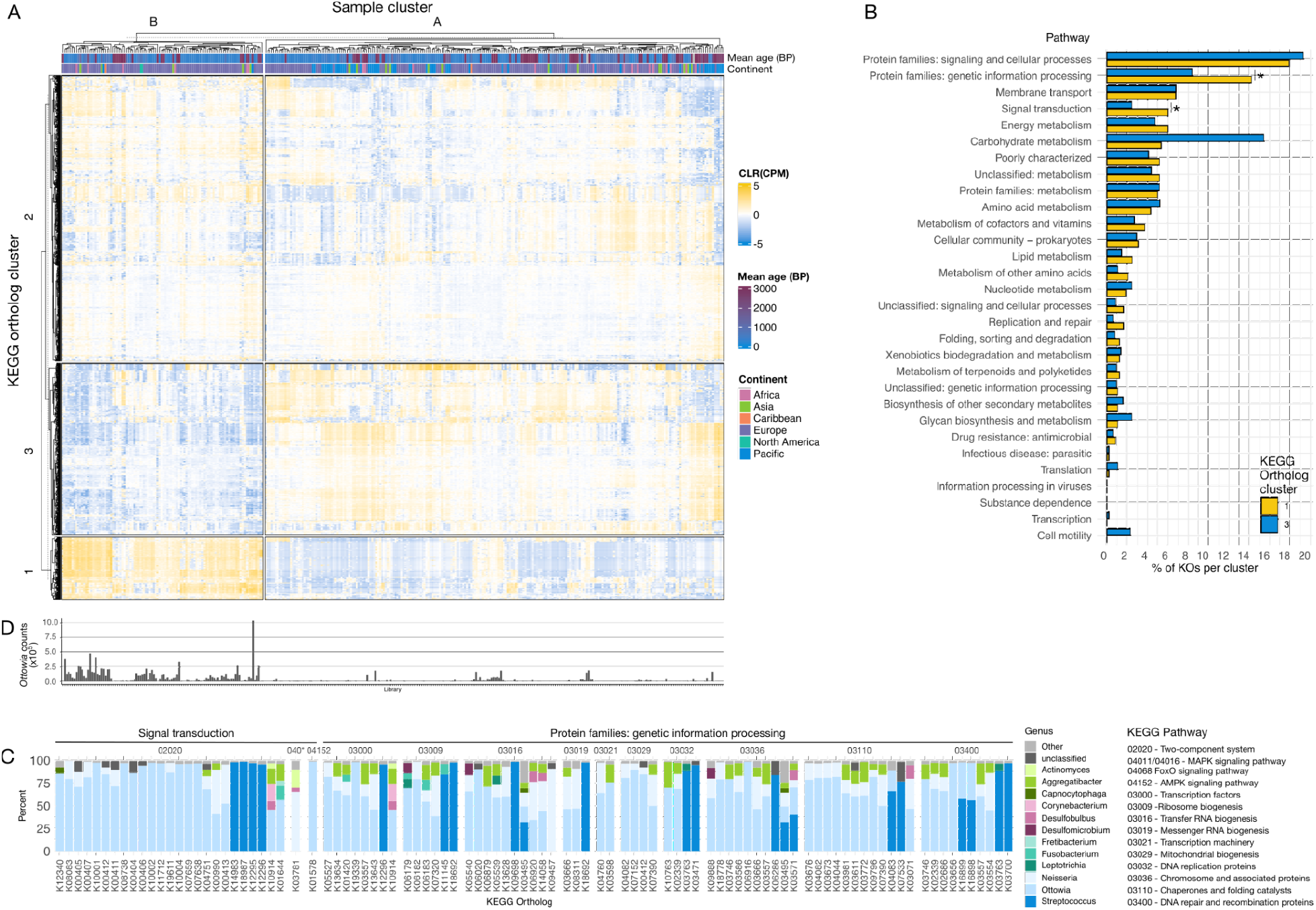
KEGG ortholog (KO) enrichment is associated with sample species composition. (**A**) Heatmap of clustered samples (clusters A and B) and KOs (clusters 1, 2, and 3), showing CLR-transformed copies per million (CPM) for each ortholog. (**B**) Percent of KOs in each KO cluster from (A) in KEGG pathways present in all samples. * p < 0.05. (**C**) Mean percent contribution by genera of KOs enriched in KO Cluster 1, which are enriched in sample Cluster B. *Ottowia* is the most prevalent genus contributing these KOs. (**D**) Read counts of *Ottowia* in all samples, showing a higher percentage of samples in cluster B have higher *Ottowia* read counts than in cluster A.

The KOs formed three clusters (Clusters 1, 2, and 3), and Sample Cluster B was enriched in KOs from Cluster 1 and depleted in KOs from Cluster 3. We grouped the KOs in Clusters 1 and 3 by Pathway and found that two pathways had more KOs in Cluster 1 than 3: Protein families: genetic information processing, and Signal transduction (Figure 4B). The genera that were contributing the orthologs in these pathways were assessed, and we found that a high proportion were attributed to *Ottowia* (Figure S7C), specifically *Ottowia* sp. oral taxon 894, a poorly characterized species. The samples in Sample Cluster B have on average higher proportions of *Ottowia* sp. oral taxon 894 than those in Sample Cluster A (Figure 4C), perhaps indicating a difference in the biofilm environment of these two clusters. Overall, *Ottowia* is the most prevalent genus enriched in KO Cluster 1 that is contributing sample Cluster B (Figure 4D).

### Phylogenetic analyses

Phylogenetic trees were constructed for *Tannerella forsythia* and *Anaerolineaceae* bacterium oral taxon 439, as both have previously been studied phylogenetically in archaeological dental calculus^16–18^. In both phylogenetic trees, samples from the same islands generally cluster together (Figure 5). The percentage of heterozygous SNPs is generally < 20% for *T. forsythia*, indicating that the reference strain used for mapping is closely related to the strains present in the samples. For *Anaerolineaceae* bacterium oral taxon 439, however, the number of sites with heterozygous SNPs is much higher, indicating that the reference genome may be quite distinct from the strains present in the samples, and reads from several strains or species in each sample may be aligning to this reference genome. Within a cluster of samples, there is a tendency for samples with higher levels of heterozygosity to fall basal to other samples (e.g., the cluster of samples from Taumako and Rurutu). It is likely that the *Anaerolineaceae* bacterium oral taxon 439 phylogeny is not that of a single taxon, but rather a collection of closely related species or strains; however, at present only one reference genome is available for oral bacteria in the family *Anaerolineaceae*.

**Figure 5.**
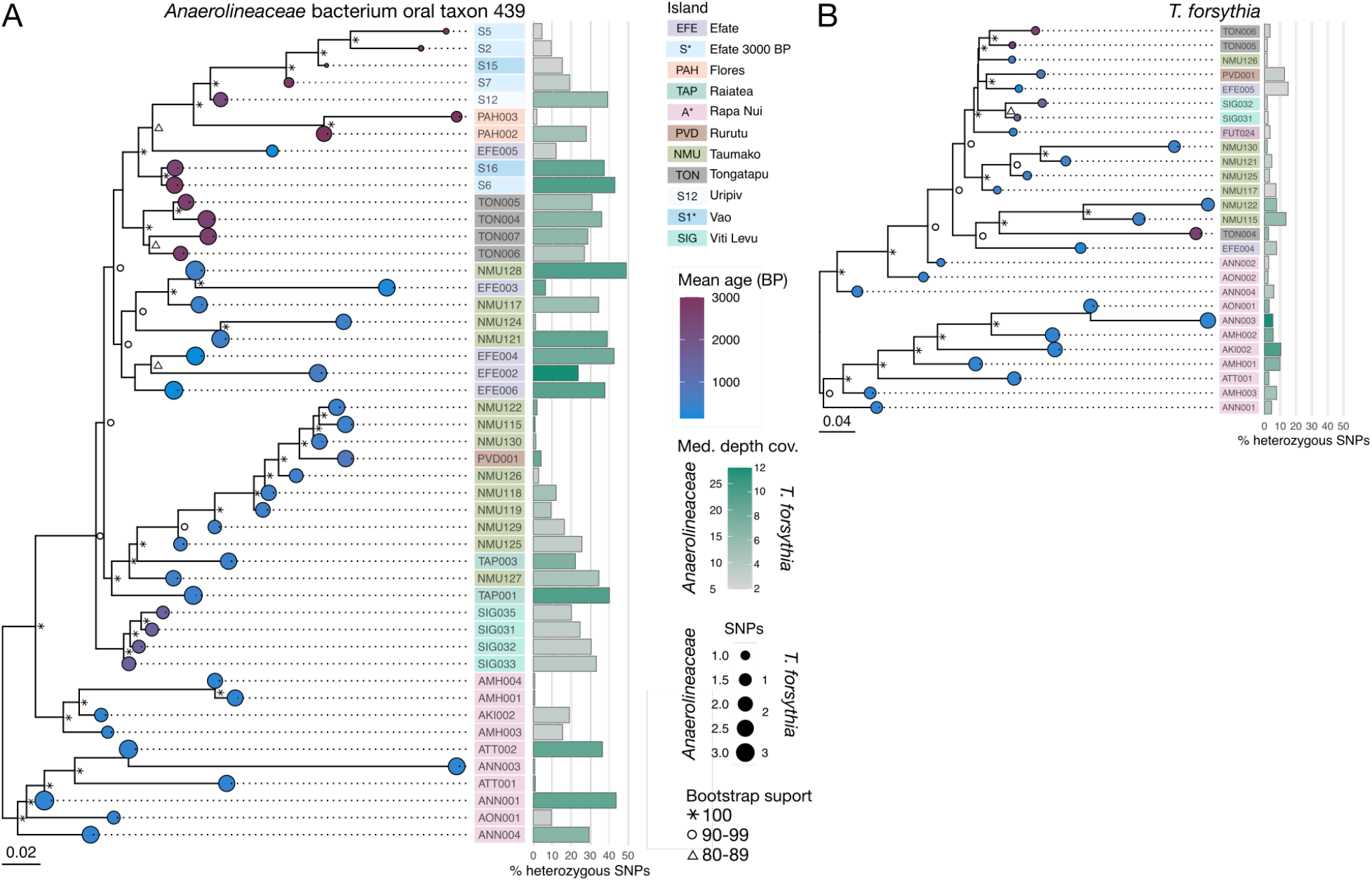
Phylogenetic trees show that bacterial genomes from the same island resemble each other. (**A**) A neighbor-joining tree of *Anaerolineaceae* bacterium oral taxon 439, including only samples with >5X genomic coverage of the taxon and using only homozygous SNPs, with midpoint rooting. (**B**) A neighbor-joining tree of *Tannerella forsythia* from samples with >2X genomic coverage, using only homozygous SNPs with midpoint rooting. For both trees, the age of the sample (in years BP) is shown as colored circles on tree tips, the circle diameter indicates the number of SNPs in that sample, the island of origin is indicated by a colored box behind sample IDs, the percentage of heterozygous SNPs is shown as a bar, and the mean coverage of the genome as the color of the bar. Scale bar indicates the genetic distance.

We additionally ran inStrain to test the ability to identify shared strains of *Anaerolineaceae* bacterium oral taxon 439 and *T. forsythia* within our samples. Following testing of inStrain with *in silico*-generated datasets to understand how ancient DNA damage patterns affect the strain assessments (Supplemental Figure S8,9), we found identical strains in samples AMH001 and AMH004 (popANI > 99.999). All other samples had popANI values <= 99.99, indicating shared, but not identical, strains (Supplemental Figure S10). No identical *T. forsythia* strains were found between any samples (Supplemental Figure S11).

As an additional test for whether there are multiple closely-related species or strains of *Anaerolineaceae* bacterium oral taxon 439 in our samples, we calculated the polymorphic rate over protein-coding genes^19^ for the reference genomes of *Anaerolineaceae* bacterium oral taxon 439 and *Tannerella forsythia*. The dN/dS values of samples mapped against the *Anaerolineaceae* bacterium oral taxon 439 genome are on average lower than for *T. forsythia*, and fall below the estimated value for mapping to an incorrect reference genome (Supplemental Figures S12, S13, indicating multiple strains of this species are likely present, while this is less likely the case for *T. forsythia*.

### Dietary DNA

In addition to tracing the microbial changes in dental calculus across the Pacific islands, we sought to examine the potential of recovering eukaryotic, food-derived DNA that may offer insight into dietary patterns across these sites. We were unable to identify any clear positive evidence of dietary DNA in our samples, which is partially due to the very low number of eukaryotic DNA sequences that were recovered, as well as trace levels of contamination and likely misalignments inaccurate reference genomes^20^ (Supplemental Table S6).

### Microparticles

To gain further insights into potential dietary patterns in the Pacific, we examined the microparticle content of the dental calculus samples analyzed in this study (Supplementary Figures S14-S18 and Supplemental Table S7). The microparticle results from Rapa Nui ^21,22^ and Teouma ^23^ were previously published elsewhere. Overall, the samples were microparticle poor compared to other dental calculus examined in the Pacific. Almost all samples contained fungal spores and hyphae, likely from sediment. There was little evidence of starch granules in the samples, and some were likely manufacturing contaminants from gloves and laboratory consumables. Low levels of phytoliths and diatoms of dietary origin were also present. They may indicate a greater reliance on starchy root crops than in Teouma, and more refined processing of root crops and better freshwater access than in Rapa Nui. However, dietary microparticles were too sparse to draw further conclusions.

## Discussion

Here we show that archaeological dental calculus samples from islands across the Pacific preserve a high proportion of DNA from endogenous oral microbiota, despite a climate that is unfavorable to DNA preservation. This allowed us to assess the diversity of ancient oral microbiomes in an understudied region, to put them in context on a global scale, and to explore the potential of the oral microbiome for tracing human migration through the Pacific. While we did not observe temporal or geographic patterns of microbiome species composition in samples from across the Pacific or across the globe, we observed a distinct community structure in the Pacific compared to samples from Europe spanning a similar time period, suggesting the presence of undescribed microbial diversity in Pacific dental calculus oral microbiomes.

Considering the high temperatures and humidity in the Pacific, and variable success in human DNA extraction from skeletal elements from the region^1^, the overall high preservation of DNA in dental calculus from the Pacific was an unexpected success. These results provide further support to the exceptional preservation of biomolecules in dental calculus^11^ compared to bones and teeth. We found that none of the climatic variables we tested predicted preservation, which suggests that smaller-scale local factors, such as soil biogeochemistry, water exposure, or the microclimate of the burial site, may be more influential. Further studies are needed to investigate local factors that may contribute to preservation, which could help explain why calculus preservation is high even in regions with conditions unfavorable to DNA preservation in skeletal remains.

Given the high level of DNA preservation in ancient dental calculus in the Pacific, we sought to explore the potential of calculus microbial DNA for tracking human migration or other behavioral or cultural changes associated with island colonization. Investigating correlations of host genetic background with community composition or strain sharing, such as by using estimated ancestry proportions for each individual, was not possible in this study, as we did not have enough calculus samples from individuals with paired human genetic data. This is, however, an exciting avenue to explore in future studies. We had limited success identifying differences in the microbial community composition related to island of origin or time period. This is in line with other studies to date, which have not found substantial microbial differences at the community-wide level related to geography, time period, or oral health^10,24–27^, indicating that the species community composition is relatively stable throughout human history. This is supported by studies of modern oral microbiomes, which indicate that perturbations in the community of dental plaque microbiomes are quickly rebalanced^28^ and are generally stable across a variety of cultural practices^29–32^, even when gut microbiomes from the same communities under comparison are substantially different.

However, there is growing evidence that the oral species present in ancient dental calculus are not fully represented in current genomic databases, such as NCBI, used for taxonomic profiling. Because of this, community composition analyses may be missing taxa and underestimating diversity, and therefore signals of differences or change may be hidden. The Pacific calculus dataset appears to be particularly affected by taxonomic database bias, as it has a notably lower taxonomic assignment rate compared to European ancient dental calculus samples for which the distributions of read length and GC content overlap.

Despite the lack of signal at the microbial community level, several studies have demonstrated the utility of phylogenetic reconstruction of abundant species in dental calculus to trace their evolutionary history, including *Anaerolinaeceae* bacterium oral taxon 439 and *Tannerella forsythia*^10,16–18,33^. Our own phylogenetic reconstructions of these species did not show clear temporal or latitudinal/longitudinal patterning; however, the reconstructed genomes often clustered by island, suggesting that oral strains are more similar within an island than between islands, which was also supported by independent strain identification with inStrain. In phylogenetic trees for both *T. forsythia* and *Anaerolineaceae* bacterium oral taxon 439, the samples from Rapa Nui, the most remote island in this study, fall in a single cluster distinct from all others, potentially reflecting that it was the last place to be colonized among the islands represented in this study^34^.

An outstanding challenge to reconstructing past microbial genomes from metagenomes is distinguishing multiple closely-related species and strains within a metagenome. Attempts to study migration patterns through the microbiome, therefore, come with a degree of inherent uncertainty when attributing microbial DNA to particular species and strains. Due to issues such as contaminated (“dirty”) reference genomes^35^, which include sequences not derived from the species of interest, or incomplete databases^36^, it is possible that sequencing reads from multiple species are inappropriately aligned to a reference genome^20,35^, creating noise in the data analysis.

This issue appears to have particularly affected our reconstruction of *Anaerolinaeceae* bacterium oral taxon 439, for which we observed high rates of SNP heterozygosity. As there is only a single isolate reference genome of *Anaerolinaeceae* bacterium oral taxon 439 sequenced to date, the extent of diversity in this organism both past and present is yet unknown. The high rate of SNP heterozygosity in many samples mapped to this reference genome appeared to affect the branching pattern within clades, despite inclusion of only biallelic SNPs in the alignment used to build our tree. Future sequencing of additional isolates of this species, or reconstruction of metagenome-assembled genomes (MAGs) of this organism from deeply sequenced modern and ancient oral metagenomes ^37^, may lead to more accurate strain separation, read alignment, SNP calling, and phylogenetic tree reconstruction. As the study of migratory patterns through ancient host-associated microbiomes is still in its infancy, method development will be fundamental in order to explore the full potential of this field.

Our results indicate the high potential of dental calculus to be well-preserved in geographic and climatic conditions that are otherwise unfavorable to DNA preservation, opening the possibility to explore archaeogenetics data in formerly poorly-accessible locations. Although we did not observe the microbial community composition of the calculus microbiome structuring by island or time period, the low taxonomic assignment rate suggests that there is additional taxonomic diversity in the Pacific calculus samples beyond that currently represented in databases, highlighting the need for studies of dental plaque biodiversity in broad, global contexts.

Individual species reconstructions have the potential to reveal evolutionary patterns that mirror the migration patterns of their human hosts, but further work disentangling closely-related species and strains within ancient dental calculus metagenomes, as well as revealing currently unknown species diversity, is needed to allow accurate identification of individual species and to perform reliable phylogenetic reconstructions.

## Materials and methods

### Laboratory methods

A total of 103 archaeological dental calculus samples were processed in this study (Table 1, Supplemental Table S1), in three labs: Jena, Oklahoma and Otago. Each lab used a slightly different extraction and library preparation protocol, detailed in the supplement. Samples from Efate were analyzed in two groups - Efate (processed in Jena, <1000 BP) and Efate 3000 BP (processed in Otago, 3000-2400 BP). Temporal information for the dental calculus samples in this study were obtained either through direct radiocarbon dating of the individual or by cultural association of the burial. Sample extraction and library preparation followed Dabney, et al.^38^, with slight variations between labs. For details see Supplementary Methods.

### General data processing

Data analyses were conducted in R v.4.1.0^39^, unless otherwise stated. General packages used were *tidyverse* v.1.3.1^40^, *readxl* v.1.3.1^41^, *ggpubr* v.0.4.0^42^ and *janitor* v.2.1.0^43^. The color palette for the study is from the R package *microshades* v.0.0.0.9000^44^. Regression models were drawn to data with a generalized linear model with geom_smooth in ggplot2 as part of tidyverse.

### Preprocessing

DNA sequencing data was preprocessed using the nf-core/eager v.2.3.3 pipeline^45^. Default options were used unless otherwise stated. Taxonomic profiling was done with MALT v.0.4.1^46,47^ with a custom database^10^.

Comparative datasets of published microbiome studies were also processed using the same procedures. One comparative dataset was used to assess preservation with the program SourceTracker^13^, and consisted of 10 non-industrialized gut samples^48,49^, 10 industrialized gut samples^50,51^, 10 skin samples^52^, 10 subgingival and 10 supragingival plaque samples^50^, 10 archaeological bone samples^10^, 10 modern dental calculus samples^10^ and 10 archaeological sediment samples^53^. In addition, 10 archaeological bone samples from Taumako and 10 from Viti Levu were included as local environmental controls; this data had been produced during genetic screening of human remains for human population genetic studies at MPI-EVA laboratories in Jena, Germany. Because bones are free of microbes during life, microbes detected in these samples provide a good proxy for local post-mortem colonization^54^.

A second comparative dataset was used to compare calculus species profiles across the globe, and consisted of ancient calculus samples from 6 previously published studies. These included samples from Japan^16^, Europe and Africa^10^, Europe^15,18,24^, and Europe, the Caribbean, North America, and Mongolia^11^ (Supplemental Table S2).

### Preservation

Preservation was assessed using SourceTracker v.1.01^13^, PCA, and the R package cuperdec^10,55^. Putative environmental and laboratory contaminants in the dental calculus samples were identified using the R package *decontam* v.1.6.0^56^, with the prevalence method. To evaluate whether preservation was related to environmental conditions, a dataset consisting of annual average temperature and annual total rainfall was compiled for each island in this study^57^. Missing data for some islands was obtained from alternative sources^58,59^. Preservation of microbial DNA in calculus and of human DNA in bone samples was compared in the Pacific and in Europe using previously published datasets^1,5,15,24,60,61^. For details see Supplemental Methods.

### Taxonomic profiling

Taxonomic profiles were generated with three tools, MALT, Kraken2, and MetaPhlAn3. The MALT table was generated as described above as part of the nf-core/eager run, and was used for all community composition analyses. Additional taxonomic profiles were generated with Kraken2 and MetaPhlAn3 to assess whether altering parameters or using non-standard databases increased the number of taxonomic assignments in samples (Supplemental Figure S3, Supplemental Figure S4). For details see Supplemental Methods.

### Community composition

Principal component analysis (PCA) was conducted on the decontaminated species table of only the Pacific data, and again on a decontaminated species table of the Pacific data plus the global comparative data, using the R package MixOmics (Supplemental Table S3). Drivers of variation in the community composition were tested with a PERMANOVA from the R-package vegan v.2.5.7^62^, with euclidean distances and 999 permutations. Homogeneity of multivariate dispersions was tested with the function betadisper from the R package vegan. Canonical correlation was performed using the R package variancePartition^63^. To determine if particular species were associated with environmental conditions on the islands, geographic location, sample age, or library characteristics, we used the R package specificity^64^ and the decontaminated species table. Pacific calculus sample clustering was performed following Quagliariello, et al.^27^, with optimal sample cluster number determined using the function clusGap in the R package cluster^65^, however the optimum number of clusters was determined to be 1 and no further cluster analysis was performed (Supplemental Figure S2). For details see Supplemental Methods.

### Functional analysis

Functional analysis was performed using HUMAnN3^66^. All reads < 50bp were removed from the fastq files prior to running HUMAnN3, because these are generally too short to be classified after translation. The bowtie2 mapping parameters were adjusted to account for aDNA damage patterns (-D 20 -R 3 -N 1 -L 20 -i S,1,0.50). The standard gene family output table with UniRef90 gene clusters was grouped to KEGG Orthology, and analysis was performed on the KEGG orthologs. Poorly preserved samples as well as those with < 750 orthologs were removed, and orthologs present at < 0.005% abundance were filtered out prior to analysis. A PCA was performed using the same steps as with taxonomy. PERMANOVA was performed with the adonis function in the R package vegan^67^.

### Phylogenetic analyses

The nf-core/eager pipeline was used, as described above, to map the non-human reads of the well-preserved samples to the abundant and prevalent oral bacteria *Tannerella forsythia* (strain 92A2, assembly GCA_00238215.1) and *Anaerolineaceae* bacterium oral taxon 439 (assembly GCA_001717545.1). Through nf-core/eager, duplicates were removed using Picard MarkDuplicates v.2.22.9, and prior to mapping, damage was clipped off of the reads (two bases for libraries with partial UDG treatment, and seven bases for non-treated libraries). Genotyping was performed with GATK UnifiedGenotyper, allowing for heterozygous calls and using all sites, with the SNP likelihood model. A minimum base coverage of 5 was required. The SNPs were further filtered in order to construct the phylogenies with only homozygous SNPs (defined as the major nucleotide having a frequency greater than 0.9), using MultiVCFanalyzer v.0.0.87^68^ (Supplemental Tables S4, S5). Only samples with at least 1000 SNPs and a mean genome-wide coverage of at least 2X (for *T. forsythia*) or 5X (*Anaerolineaceae* bacterium oral taxon 439) were included. The coverage requirement was increased for *Anaerolineaceae* bacterium oral taxon 439, because its percentage of heterozygous SNPs was higher. Neighbor-joining trees were built using distance matrices generated with the TN93+G4 substitution model, as this was determined by testing with the DECIPHER R package^69,70^ to be the best model for each dataset. The trees were rooted using the midpoint, determined with the package R phangorn^71^. The phylogenetic trees were constructed and visualized using R packages ape v.5.5^72^, ade4 v.1.7.17^73^, adegenet v.2.1.3^74^ and ggtree v.1.99.1^75^.

### Strain sharing

Strain sharing across samples was assessed with inStrain^76^. A test dataset was generated *in silico* to test for the effects of ancient DNA damage on popANI calculations. For details see Supplemental Methods. The coverage overlap and popANI between all samples was visualized and compared in R (Supplemental Figures S8 and S9). Strain sharing across Pacific calculus samples was assessed with inStrain for 6 species that were highly abundant across the Pacific calculus dataset: *Actinomyces dentalis* (GCF_000429225.1), *Anaerolineaceae* bacterium oral taxon 439 (CP017039.1), *Desulfobulbus oralis* (CP021255.1), *Eubacterium minutum* (CP016202.1), *Olsenella* oral taxon 807 (CP012069.2), and *Tannerella forsythia* (NC_016610.1). We focused only on *Anaerolineaceae* bacterium oral taxon 439 and *Tannerella forsythia*, for which we generated whole genome SNP-based phylogenies.

The script polymut.py from cmseq^19^ was used to assess whether samples contained multiple strains of the species *Tannerella forsythia* and *Anaerolineaceae* bacterium oral taxon 439 based on the ratio of non-synonymous vs. synonymous sites (dN/dS). We performed a test to determine the expected dN/dS of mapping reads against an incorrect reference genome by using three species of *Fusoboacterium*, which were formerly considered subspecies of *F. nucleatum*: *F. nucleatum*, *F. polymorphum*, and *F. vincentii*. We calculated the ANI of each genome compared to the other two (Supplemental Figure S12A) and used polymut.py to calculate the dN/dS for short read datasets of each genome mapped to the three reference genomes (Supplemental Figure S12B-G), and took the average across all genomes, which was 1.94. We took this to be the expected dN/dS value when mapping a species against a closely-related but incorrect reference genome. For our samples, collapsed reads were mapped against the *Tannerella forsythia* genome (NC_016610.1) and the *Anaerolineaceae* bacterium oral taxon 439 genome (CP017039.1) using bwa aln and the flags -n 0.01 -l 1024 to account for ancient DNA damage. We then calculated the ratio of non-synonymous SNPs to synonymous SNPs (dN/dS) for all samples mapped against the *Anaerolineaceae* bacterium oral taxon 439 or *Tannerella forsythia* genome (Supplemental Figure S13).

### Eukaryotic DNA

In addition to analyzing microbial DNA, we also investigated putative eukaryotic DNA within the samples. nf-core/eager was used to align the DNA sequences to reference genomes (Supplemental Table S6) for the five species of interest (with mapping quality set to 37): cattle (*Bos taurus*), dog (*Canis lupus familiaris*), broad fish tapeworm (*Dibothriocephalus latus*), bamboo (*Fargesia denudata*), and wheat (*Triticum aestivum*). Damage patterns were investigated for individuals with at least 200 reads mapping to the specific taxa^20^. For taxa where 200 reads were not reached for any individuals, the damage patterns of the 10 individuals with the highest number of reads were investigated.

### Microfossil analysis

Methods used for Rapa Nui^21,22^ and Teouma ^23^ were previously published. All other samples had been decalcified in 0.5 M EDTA for aDNA extraction. After aDNA extraction, the remaining EDTA aliquot was rinsed in DDI water and a 40 μl drop containing most of the microparticle sample was placed on a glass slide and covered with a glass coverslip. Optical light microscopy was done with a Zeiss Axioscope (located in the Anthropology and Archaeology Department at the University of Otago). Each slide was examined in its entirety in vertical transects using cross-polarized and transmitted light to identify and photograph all microparticles. All microparticles were counted and separated into plant or animal types and then by morphotype. Identification of microparticles was done using published material as well as the Pacific-focused reference collection developed by MT. The International Code for Phytolith Nomenclature 2.0 was used to name and describe all phytoliths ^77^.

## Data Accessibility

The sequencing data for this study is deposited in the European Nucleotide Archive under accession PRJEB61887. The scripts used for analysis and figure generation are available on github at https://github.com/ivelsko/pacific_calculus.

## Acknowledgements

We thank R. Shing and M. Abong of the Vanuatu Cultural Centre (VCC) for their assistance during calculus sampling. Sampling of the Teouma dental calculus was done through a research agreement between M.T. and the Vanuatu National Cultural Council. The Teouma Archaeological Project is a joint initiative of the Vanuatu National Museum and the Australian National University (ANU), directed by M.S. and S.B. and at different times R. Regenvanu and M. Abong, both former Directors of the VCC. Funding for the project was provided by the Australian Research Council (grant no. DP 0556874), the National Geographic Society (grant no. SRC 8038–06), the Pacific Biological Foundation, the Department of Archaeology and Natural History and School of Archaeology and Anthropology at the ANU, the Snowy Mountains Engineering Corporation Foundation and B. Powell. The laboratory research and travel for excavation of the skeletal remains were funded by The Royal Society of New Zealand Marsden Fund (grant nos. UOO0407 and 09-UOO-106) and a University of Otago Research Grant awarded to H.B. The support of the leaseholder M. R. Monvoisin and family is acknowledged, as is the support and assistance of the traditional landowners and population of Eratap Village. Detailed acknowledgement by the authors of the Vanuatu studies (SB, HB, JF, RS, MS, EW, and FV) are given in the published site reports for each location, and were supported by the Australian Research Council DP160103578. We thank Alexander Hübner for discussions on strain analyses. This work was supported by the Werner Siemens Stiftung (“Paleobiotechnology” to C.W.), the Deutsche Forschungsgemeinschaft (DFG, German Research Foundation) under Germany’s Excellence Strategy EXC 2051, Project-ID 390713860 (C.W.), and the Max Planck Society. The Otago research presented here was funded by a University of Otago Doctoral Scholarship, a Royal Society of New Zealand Skinner Fund grant and an Otago Centre for Electron Microscopy Student Research Award awarded to M.T.

## Author contributions

I.M.V., Z.F., and M.T. performed data processing, analysis, interpretation, and wrote the manuscript. Z.F. and M.T. performed data generation. S.B., H.R.B., G.C., J.D., J.F., A. L.-T., C.M. L., E. M.-S., K. N., A.T.O., C.P., R.S., M.S., E.W., and F.V. provided samples, resources, and archaeological information. A.B.R. provided resources. C.W. designed the study and wrote the manuscript.

## Supplemental Methods

### Laboratory methods

#### Jena

All sample processing took place in a dedicated cleanroom facility at the Max Planck Institute for Evolutionary Anthropology laboratories (Jena, Germany). Total DNA was extracted from 0.5-7 mg of dental calculus per individual, using a silica column-based extraction protocol optimized for the recovery of short DNA fragments, adapted for dental calculus ^1–3^. The extracts were prepared into double-stranded libraries with partial uracil-DNA-glycosylase (UDG) treatment ^4,5^ and dual indexing ^6–8^. The libraries were sequenced to a depth of 10.5 ± 2.3 million reads (mean ± standard deviation) on an Illumina Nextseq with 75-bp paired-end sequencing chemistry. Blanks were processed alongside the samples for both extraction and library preparation. Two samples, SIG040 and SIG046, were extracted and sequenced twice, as it was suspected that burial sand may have been unintentionally included during the first processing round.

#### Oklahoma

Total DNA was extracted from 0.8-12.8 mg dental calculus per individual following Ozga *et al.* ^9^, at the ancient DNA facility of the University of Oklahoma Laboratories of Molecular Anthropology and Microbiome Research (LMAMR, Norman, OK, USA). This method is very similar to that performed in Jena. Blanks were processed alongside the samples. The extracts were thereafter shipped to MPI-EVA, where they were prepared into libraries, as described above, alongside the Jena samples. The samples were sequenced to a depth of 10.6 ± 1.5 million reads on the same NextSeq flow cells as the Jena samples with 75-bp paired-end sequencing chemistry.

#### Otago

Total DNA was extracted from approximately 1-17 mg of dental calculus per individual using a phenol-chloroform aDNA extraction protocol ^10^. Dental calculus samples were washed with ultrapure water and allowed to dry in a laminar flow hood overnight. A second wash was performed using 1 ml of 0.5 M EDTA, with a 30 min incubation time. The supernatant was removed, and the samples were demineralized in 1 ml of 0.5 M EDTA for up to 72 hours, until fully demineralized. The supernatant was added to a tube with 750 μl of phenol:chloroform:isoamyl (25:24:1), vortexed, and left on a rotator for 10 min. After centrifuging, the aqueous phase was transferred to 750 μl of phenol:chloroform:isoamyl alcohol (25:24:1). The incubation step was repeated, after which the aqueous phase was transferred to 750 μl of chloroform:isoamyl alcohol (24:1). After vortexing and mixing by inversion, the mixture was centrifuged and the aqueous phase transferred to 13 ml of 6 M GuSCN and 200 μl of silica suspension, and left on a nutator for 30 min. After centrifugation, the supernatant was removed, the silica was resuspended in 1 ml of GuSCN binding buffer, and the supernatant discarded after centrifugation (three times in total). The silica pellet was air dried for 15 min, and DNA eluted twice in 60 μl TE (heated to 65-75°C). Blanks were processed alongside samples through extraction and library preparation. Double-stranded libraries were prepared by blunt-end repairing the DNA strands, and after that ligating and filling in adapters. The libraries were amplified using KAPA HiFi enzyme, and no UDG-treatment was performed. The libraries were sequenced using an Illumina MiSeq 75-bp paired-end sequencing chemistry to 8.3 ± 7.2 million reads at the Otago Genomics Facility (Otago, New Zealand).

### Data processing

Within the nf-core/eager pipeline, poly-G stretches were removed from the raw data, as they are a common by-product of the two-color chemistry sequencing strategy used by Illumina’s NextSeq. Human DNA was removed from the dataset by mapping to the human reference genome GRCh38, and only unmapped reads were retained for downstream microbiome analyses. To produce an OTU table of the microbes present in the samples, the dataset was aligned using MALT v.0.4.1 ^11,12^ to a custom database containing all bacterial and archaeal assemblies (scaffold/chromosome/complete levels, up to 10 randomly selected genomes per species) from RefSeq and the human HG19 reference genome ^13^ via the nf-core/eager pipeline. In addition, the dataset was aligned to the NCBI nt database (as of October 2017), to screen for eukaryotic DNA. MEGAN v.6.17.0 ^14^ was used to export OTU tables from the resulting MALT-produced rma6 files, using summarized read counts at both the genus and species level.

### Preservation

Preservation was investigated and visualized using SourceTracker v.1.01 ^15^. A species-level OTU table was used as input, and the reference metagenomic datasets described in the main text were used as sources. During the SourceTracker analysis, rarefaction was performed to 10,000 reads, with a training data rarefaction of 5,000 reads. For principal component analysis (PCA), species-level read counts of all dental calculus samples and sources (including an additional 9 modern dental calculus samples from the same study as listed in the main text) were compared. The R-package cuperdec ^13,16^ was used to identify well-preserved samples (adaptive burn-in method, cut-off 50%), and 73 (out of 103) samples were carried on to further analyses based on this analysis.

To remove potential contaminant species, we used the R package decontam. The samples were separated into groups based on the processing lab, and blanks and archaeological bones were used as proxies for contamination sources (cut-off 0.25 for all groups, for both blanks and bones). Contaminants from all labs were combined into a single list of all contaminants, and all contaminants were removed from all samples. The comparative calculus datasets were likewise assessed with decontam and all contaminants were combined into a single list, along with those from the Pacific calculus dataset here, and all contaminants were removed from all datasets for the comparative analyses.

Annual average evapotranspiration was compiled from 1717. The proportion of taxa estimated to originate from the oral microbiome (i.e., dental plaque and dental calculus) by SourceTracker was used as a proxy for preservation of the archaeological dental calculus samples. The effects of environmental variables and/or sample age on preservation were investigated using beta regression with a complementary log-log link function to account for observed heteroscedasticity. Using ANOVA, it was found that the model was not significantly improved by adding the random effects of laboratory and/or island (p»1). Step AIC (using both directions) was used for model selection. To reliably estimate parameters for the model, statistically influential data points were removed, and the model-fitting process was repeated until a stable dataset was reached. Model fit was measured using the R^2^ value as suggested by Ferrari and Cribari-Neto ^18^ for beta regression models.

Preservation of calculus was determined by the percentage of the sample that was determined to be from an oral source (calculus, supragingival plaque, and subgingival plaque) with SourceTracker. Preservation of human DNA was determined by the percentage of endogenous human DNA in shotgun sequenced data, and was taken from published values. Human DNA preservation in the Pacific was taken from Posth, et al. 2018 (Supplemental Table S3) ^19^ and Liu, et al. 2022 (Supplemental Table S2) ^20^, human preservation in England was from Schiffels, et al. 2016 (Supplemental Table S1) ^21^ and Patterson, et al. 2022 (Supplemental Table S1) ^22^, and calculus preservation in Europe was taken from SourceTracker values of data from ^23,24^.

### Taxonomic profiling

Kraken2 was run with default parameters, using two databases (described in ^25^): a custom RefSeq database and the same custom RefSeq database with the addition of MAGs from Pasolli, et al. (2019). This allowed us to test whether including additional diversity in the database resulted in a substantial increase in the number of reads assigned to taxonomy. However, we found that it did not, and that this profiler is particularly affected by the read length. MetaPhlAn3 was run with two different sets of parameters: default, and custom settings optimized for assignment of ancient DNA (-D 20 -R 3 -N 1 -L 20 -i S,1,0.50 and minimum read length of 35 bp). The use of ancient-optimized parameters substantially increased the number of species identified in each sample (Supplemental Figure S4A) and we found that the number of species identified by MetaPhlAn3 with ancient-optimized parameters and by MALT were correlated (Supplemental Figure S4B). The optimal number of sample clusters within the Pacific calculus dataset was determined using the Gap statistic with clusGap from the R package cluster^26^ with 500 bootstrap replicates, on both the same MALT species table used for compositional analysis, as well as on a MALT genus table that was cleaned and filtered following the same steps as the species table. The optimal number of clusters was found to be one for both the species and genus tables (Supplemental Figure S2), so no further cluster analysis was performed.

### Strain sharing

Simulated metagenome testing. Strain sharing across samples was assessed with inStrain ^27^. Two 10M bp paired-end simulated read datasets were generated to test the effects of aDNA damage patterns and clipping, using the genomes of the 10 species in the ZymoBiomics kit that was used by Olm, et al. for testing, in the proportions described in the kit. The genomes were processed with gargammel ^28^ in two batches, to create datasets with two different damage profiles based on the damage of the samples sequenced here. Two samples were selected for references, HCLVMBCX2-3505-07-00-01_S7, which is non-UDG treated, and EFE002.B0101, which is UDG-half treated. Both samples were mapped against the *Anaerolineaceae* bacterium oral taxon 439 genome (GCA_001717545.1) with bwa (-n 0.02 -l 1024) ^29^, and the bam files were used as input to mapDamage ^30^ to generate damage profiles and read length distributions that were used as input to gargammel. The simulated datasets were processed with the nf-core/eager pipeline the same way as samples, and adapter-trimmed un-collapsed paired reads were pulled out of the eager output with a custom script. The paired end reads were mapped against all 10 reference genomes in the ZymoBiomics kit, combined into a single fasta file with bwa (-n 0.02 -l 1024). The effect of read end masking to remove aDNA damage was tested with a custom python script ^31^, with mapped reads from UDG-treated reads masked 1 base on either end, and non-UDG-treated reads masked 9, 11, 13, and 15 bases on either end, based on the C-T transversion ratio along read lengths determined by mapDamage. InStrain profile was run on the unmasked and masked mapped read bam files with insert sizes 12, 24, 36, and 48, and inStrain compare was run on all output of the profile step. Based on the output of these simulated data tests, a mask length of 11 and insert size of 12 were selected for processing real samples, as these returned a maximum popANI with minimal loss of coverage.

Pacific calculus samples. Strain sharing across samples was assessed with inStrain for 6 species that were highly abundant across the Pacific calculus dataset. Paired-end reads were mapped against reference genomes for *Actinomyces dentalis* (GCF_000429225.1), *Anaerolineaceae* bacterium oral taxon 439 (CP017039.1), *Desulfobulbus oralis* (CP021255.1), *Eubacterium minutum* (CP016202.1), *Olsenella* oral taxon 807 (CP012069.2), and *Tannerella forsythia* (NC_016610.1), combined into a single fasta file. Only *Anaerolineaceae* bacterium oral taxon 439 and *Tannerella forsythia* had sufficient genome breadth and depth of coverage across enough samples for reliable strain comparison with inStrain. Mapped reads in each bam file were masked according to their UDG treatment, with UDG-half-treated samples masked 1 base at both ends, and non-UDG-treated samples masked 11 bases at both ends. inStrain profile was run on the masked bam files with an insert size of 12, and the default value of 5X minimum coverage was required for samples to be included in inStrain analysis. inStrain compare was run on all output files of inStrain profile. We focused only on *Anaerolineaceae* bacterium oral taxon 439 and *Tannerella forsythia*, for which we generated whole genome SNP-based phylogenies.

The script polymut from cmseq ^32^ was used to assess whether samples contained multiple strains of the species *Tannerella forsythia* and *Anaerolineaceae* bacterium oral taxon 439 based on the ratio of non-synonymous vs. synonymous sites (dN/dS). We performed a test to determine the expected dN/dS of mapping reads against an incorrect reference genome by using three species of *Fusoboacterium*, which were formerly considered subspecies of *F. nucleatum*: *F*. *nucleatum*, *F. polymorphum*, and *F. vincentii*. These genomes were run through prokka to generate gff files for polymut. The ANI of each genome compared to the other two was determined with MASH ^33^ as part of the program dRep ^34^ and found to be below the species-cutoff of 95% identity (Supplemental Figure S12A). Nine simulated short-read datasets were generated from each of the three genomes with three read lengths and three different levels of deamination: long read length (100bp), medium read length (75bp), and short read length (50bp), and no deamination, low deamination, and high deamination. These 9 short-read datasets were each mapped against all three reference genomes using bwa aln and the flags -n 0.01 -l 1024, and the script polymut.py was used to determine the number of synonymous SNPs, the number of non-synonymous SNPs, and the total number of sites compared, for each mapped bam file (Supplemental figure S12 1B-G). We found that the deamination level did not affect the dN/dS values for any of the species mapped to any of the others. Because the Pacific dataset reads have a short read length, we focused on the results of the short read length dataset, and calculated the average dN/dS for short reads across all deamination levels, which is 1.94. We took this to be the expected dN/dS value when mapping a species against a closely-related but incorrect reference genome.

For our samples, collapsed reads were mapped against the *Tannerella forsythia* genome (NC_016610.1) and the *Anaerolineaceae* bacterium oral taxon 439 genome (CP017039.1) using bwa aln and the flags -n 0.01 -l 1024. Mapped reads were masked according to their UDG treatment as described above (i.e. UDG-half treated reads were masked 1bp on both ends, and non-UDG treated reads were masked 11bp on each end). Masked mapped bam files were run through polymut with a minimum coverage requirement of 5X, min quality 30, and dominant frequency threshold of 0.8, and the gff files from NCBI RefSeq for the genome accessions above.

### Eukaryotic DNA

Identifying eukaryotic taxa within metagenomic datasets is challenging and requires multiple validation steps ^35^. Additionally, it should be noted that some of the potential dietary items, such as kava (*Piper methysticum*) and the giant swamp taro (*Cyrtosperma merkusii*), from the Pacific may not be present in reference databases as they do not have sequenced nuclear or organelle genomes. Within the well-preserved dental calculus samples, we identified DNA from five non-human eukaryotic species of interest: cattle (*Bos taurus*), dog (*Canis lupus familiaris*), broad fish tapeworm (*Dibothriocephalus latus*), bamboo (*Fargesia denudata*), and wheat (*Triticum aestivum*) (Table S1). Other eukaryotic DNA present in the dataset belonged to commonly recognized contaminants, were assigned to genomes with known contaminating sequences ^35^, or represented highly unlikely taxa; these assignments were excluded from subsequent analysis. For cattle, mapping was also performed to the zebu genome (*Bos indicus*), as this species may be a closer match for this region ^36^. For bamboo, only the chloroplast genome was available. For wheat, the mapping was restricted to the mitochondrial genome in this initial step, as the full wheat genome is very large (15.4 Gb).

### Microparticle analysis

Fifty-four samples of decalcified dental calculus were analyzed in this study – samples that were decalcified using EDTA in Jena were sent to Otago for analysis after DNA extraction. Phytoliths and starch granules were identified based on MT’s reference collection from several botanic gardens and herbariums in Vanuatu, New Zealand, Hawaii, and Rapa Nui, as well as other published material (for example ^37^). Phytoliths were described based on the International Code for Phytolith Nomenclature (ICPN) 2.0 ^38^.

Efate. The samples from Efate were not very microparticle rich; the most common microparticles were fungal spores and hyphae (commonly found in sediment samples and not possible to refine the ID) (Supplemental Figure S14A). Sample EFE002.B did contain diatoms, but since it is only one sample and two diatoms, not much can be inferred from this. One starch granule was found in sample EFE003.B; however, it is round and less than 10 µm, which means it could come from just about any starch-containing plant (Supplemental Figure S14B). A starch granule was also found in sample EFE006.B (Supplemental Figure S14C). This granule is approximately 10 µm and angular; it is most similar to *Colocasia esculenta* or taro; however, it could also overlap with several other root crops and finding it in isolation makes it difficult to be certain.

Futuna. The Futuna samples had the highest number of starch granules of any location examined, and almost all were in one sample, FUT018.B. Of the 27 granules, 18 are ≤ 10 µm and so cannot be confidently identified to family/species. The remaining 9 granules correspond to Types 2 (n=3), 3 (n=2), 4 (n=2), 5 (n=1) and 8 (n=1) (Supplemental Figure S15A,B,C,D,E) ^10^. Type 2 granules could come from several root and tree crops and corn, which could be contamination, although the possibility that it is a genuine dietary component cannot be ruled out given that Europeans may have traded corn there as early as the 1600s ^39,40^. Type 3 granules could be from several root and tree crops. Type 4 granules are generally from *Tacca leontopetaloides* or Polynesian arrowroot – unfortunately, they also overlap with corn, so contamination cannot be ruled out. The type 5 granule could be from several root and tree crops. The type 8 granule is similar to one found in Lapita-aged samples from Vanuatu but could not be linked to any known reference species. A few different fungal spores and hyphae were also found in this sample. Sample FUT021.B contained one starch granule consistent with *Zea mays* (corn), which is likely contamination. There was also a piece of microcharcoal (Supplemental Figure S15F) and three probable grass or Cyperaceae phytoliths (Supplemental Figure S15G,H,I) in this sample.

Taumako. Almost all samples from Taumako contained fungal spores and/or hyphae, but not much else. Five starch granules were found in three samples – all are either too small or damaged to be securely identified except for one, which is consistent with wheat starch and likely contamination (Supplemental Figure S16A,B, C). Sample NMU116.A contained unusual microparticles that remain unidentified (Supplemental Figure S16D); these may be associated with the high marine diet found in stable isotope results from the same population ^41^. Finally, a few phytoliths were recovered; most were dentate elongate and likely from grass leaves; some were damaged and could not be adequately described (Supplemental Figure S16E,F).

Fiji. There were quite a few phytoliths found in the Fiji samples, most of which are probably from grass leaves (blocky, elongate, bilobate and rondel types), along with a couple of samples that contained palm phytoliths (Supplemental Figure S17A-J). These are all quite commonly found within the Pacific. There was one phytolith that could be species-specific that did not match anything in our reference database (Supplemental Figure 17B); further reference samples would be needed to positively identify it, but it resembles phytoliths from the bark of species with medicinal properties in West Africa ^42^. There are also two phytoliths that look like double-peaked glume rice phytoliths (Supplemental Figure 17A,C), but they are very degraded so the resemblance may be an artifact. Several of the phytoliths appeared black or burnt (Supplemental Figure 17G,H,I), which may be an indication of fire use. Overall, the Fijian samples were very mineral rich, likely due to insufficient removal of residual sediment for these samples (this issue is also noted in the aDNA methods above, where two samples had to be re-extracted and sequenced due to sediment contamination). Several of the mineral particles are large (probably too large to be inclusions in the dental calculus) and olive green (however, they are unlikely to be obsidian due to the lack of conchoidal fractures) (Supplemental Figure S17K). Sample SIG044.A contained an unknown fiber probably of plant origin as there are no visible scales or a medulla; it may also be part of an insect (Supplemental Figure S17L).

Tongatapu. Most samples from Tongatapu contained at least one phytolith. There were mostly palm phytoliths (spheroid echinate) (Supplemental Figure 18B), followed by probable grass phytoliths (Supplemental Figure 18A) and one instance of a Cyperaceae phytolith (Supplemental Figure 18C). Two samples also contained sponge spicules (Supplemental Figure 18D). One sample, TON001.C contained a unique looking fiber that may be a taphonomically damaged feather barbule (Supplemental Figure 18E).

**Supplemental Figure S1.**
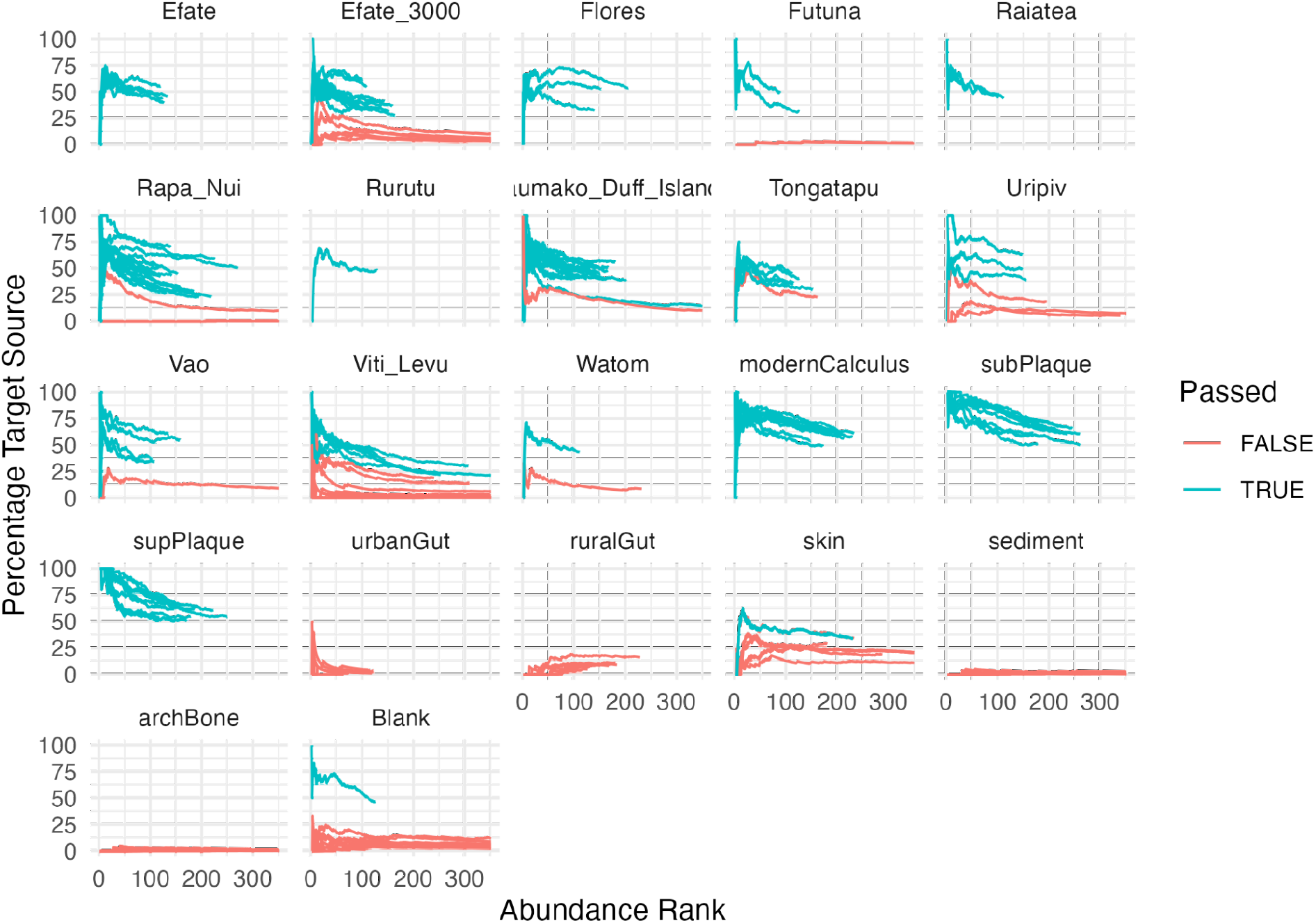
Cumulative percent decay (cuperdec) curves for newly sequenced samples from the Pacific presented in this study. Samples are grouped by island. Passed True indicates a sample passed the cut-off and is well-preserved, while Passed False indicates a sample did not pass the cut-off and is not well-preserved.

**Supplemental Figure S2.**
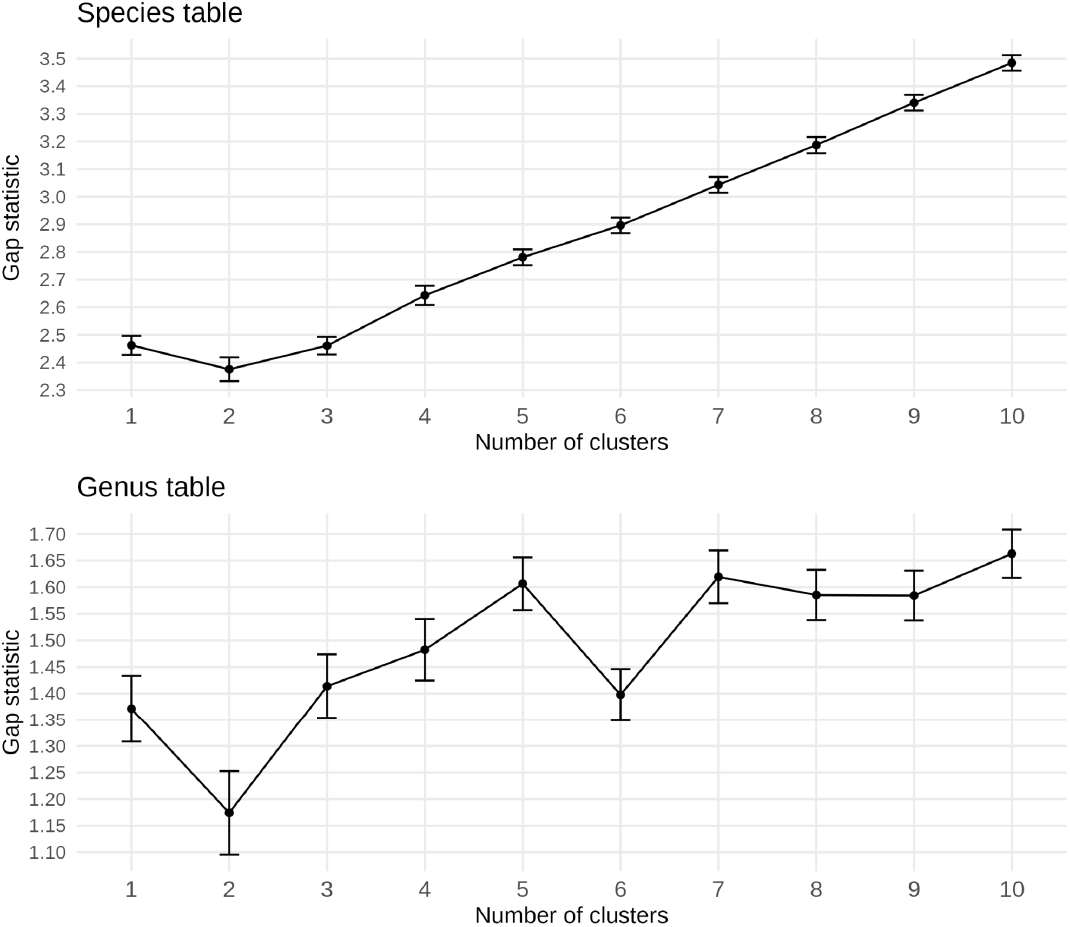
Gap statistic to test for the optimal number of sample clusters in the Pacific calculus dataset. The upper panel is based on the species table, while the lower panel is based on the genus table. The Gap statistic tests the goodness of fit of the number of clusters in a dataset. The optimal number of clusters is determined as the cluster before the first drop in value of the Gap statistic. When using either the species table or the genus table, the first drop in the Gap statistic occurs at 2 clusters, such that the optimal number of clusters in the samples is 1. Error bars indicate standard error of the mean based on 500 bootstrap replicates.

**Supplemental Figure S3.**
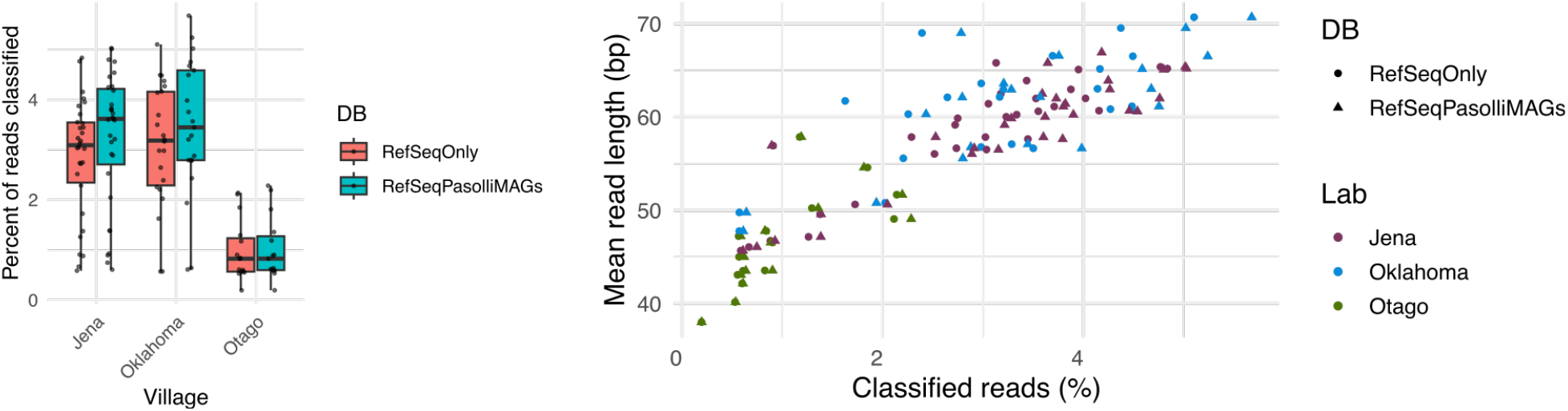
Inability to increase read taxonomic assignment by using a database with novel metagenome-assembled genomes (MAGs) not found in the NCBI RefSeq database. A. Percent of reads per sample that were assigned taxonomy when using the taxonomic profiler Kraken2 and a database consisting of RefSeq genomes only, or a database of the same RefSeq genomes plus MAGs published by Pasolli, et al. (2019). B. Correlation between the percentage of classified reads per sample and the average read length in each sample. Samples with longer average read length have higher average percent of reads assigned taxonomy.

**Supplemental Figure S4.**
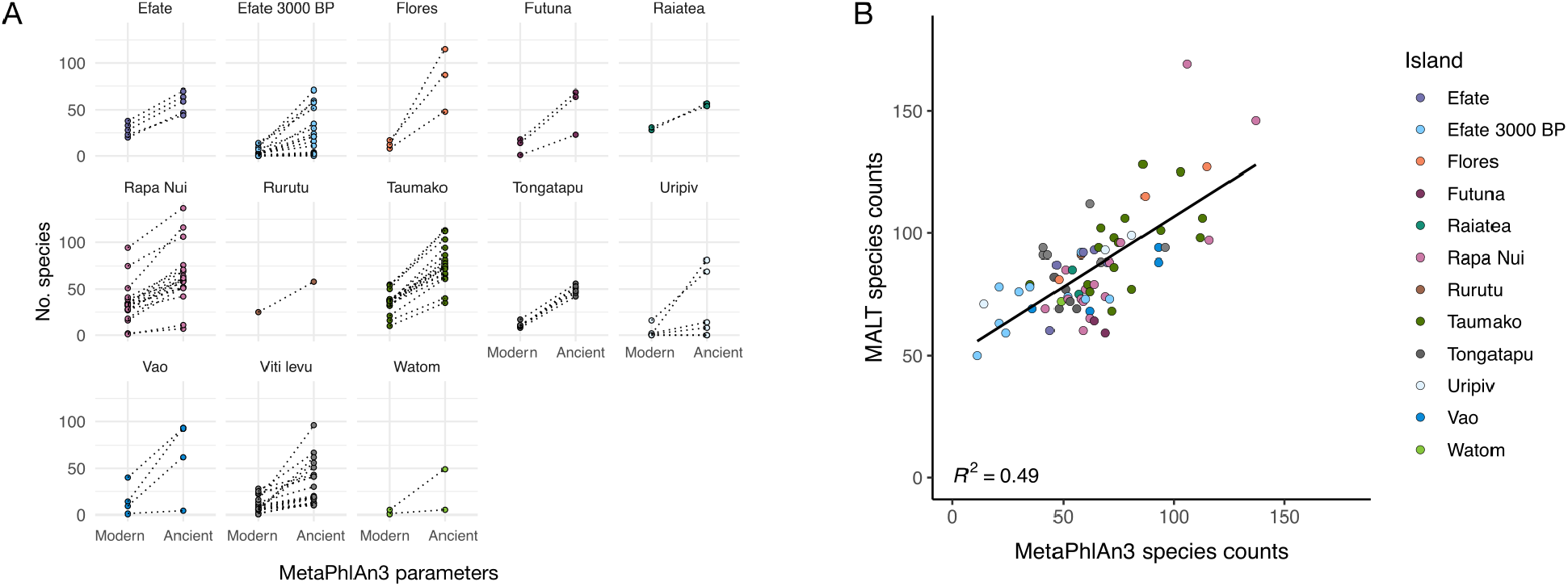
Species counts by MetaPhlAn3. A. Number of species detected per sample by MetaPhlAn3 using modern or ancient parameters for the bowtie2 mapping step. B. Concordance between the number of species detected by MALT after filtering and by MetaPhlAn3 run with ancient parameters.

**Supplemental Figure S5.**
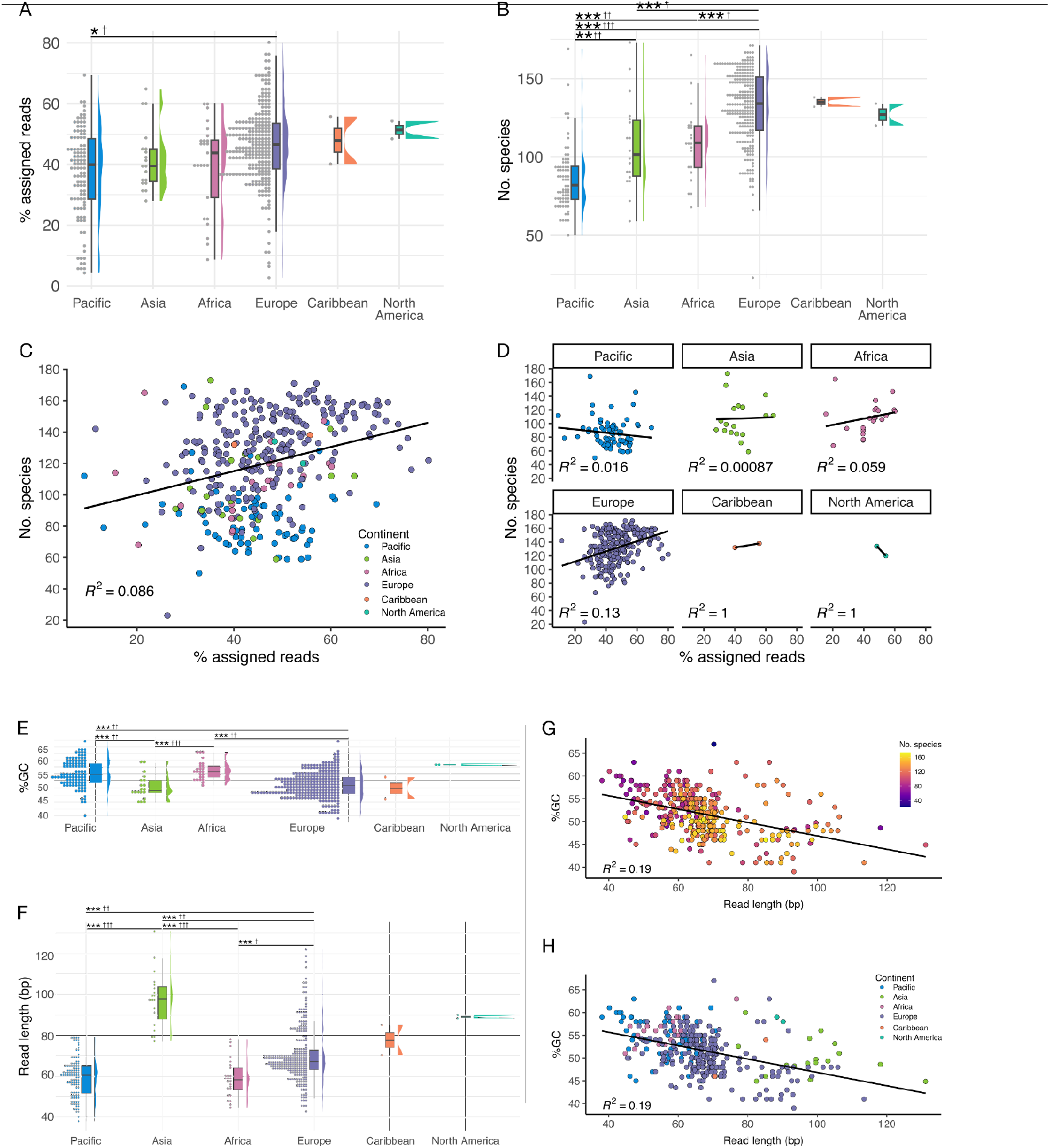
Read and species assignment rates. **A.** The percentage of reads in each sample that was assigned taxonomy by MALT, grouped by continent of origin. **B.** The number of species detected in each sample after filtering. **C.** Number of species and percent of assigned reads in samples. **D.** Same as **C** but separated by continent. The Pacific samples have a slight negative association between the percent of assigned reads and the number of detected species. **E.** Average GC content of the reads per sample, grouped by continent. **F.** Average read length per sample, grouped by continent. G. Average GC content, read length per sample, and number of species assigned per sample, are weakly negatively correlated. **H.** Same as **G** but colored by continent of origin. ** p < 0.01, *** p < 0.001, † effect size < 0.3, †† effect size >= 0.3, <= 0.5, ††† effect size > 0.5.

**Supplemental Figure S6.**
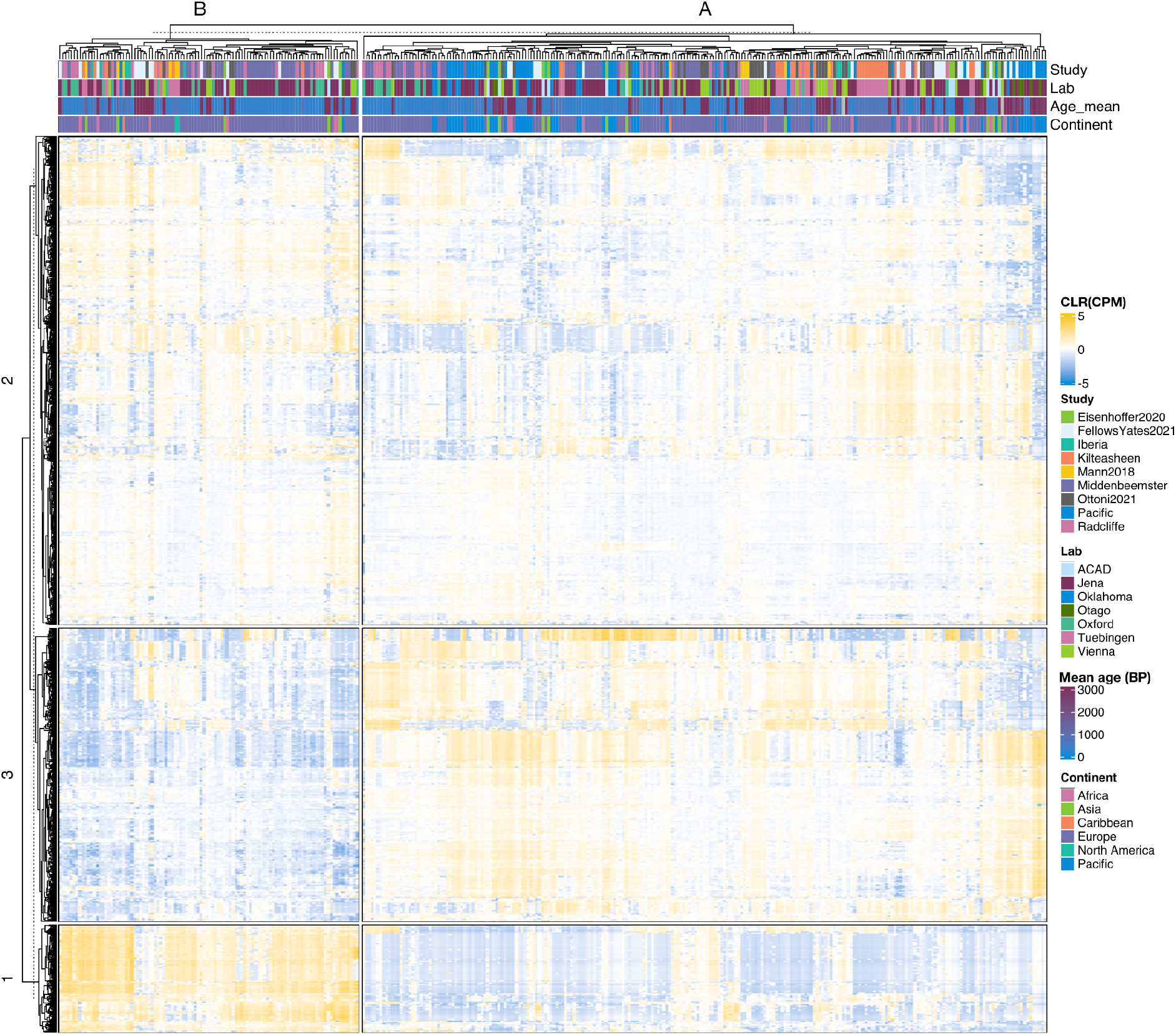
Hierarchically clustered heatmap of KEGG ortholog abundance in all samples (CLR-transformed copies per million), same as main Figure 4A, but with additional sample metadata shown across the top including the lab in which sample processing was performed, and the study in which samples were originally published.

**Supplemental Figure S7.**
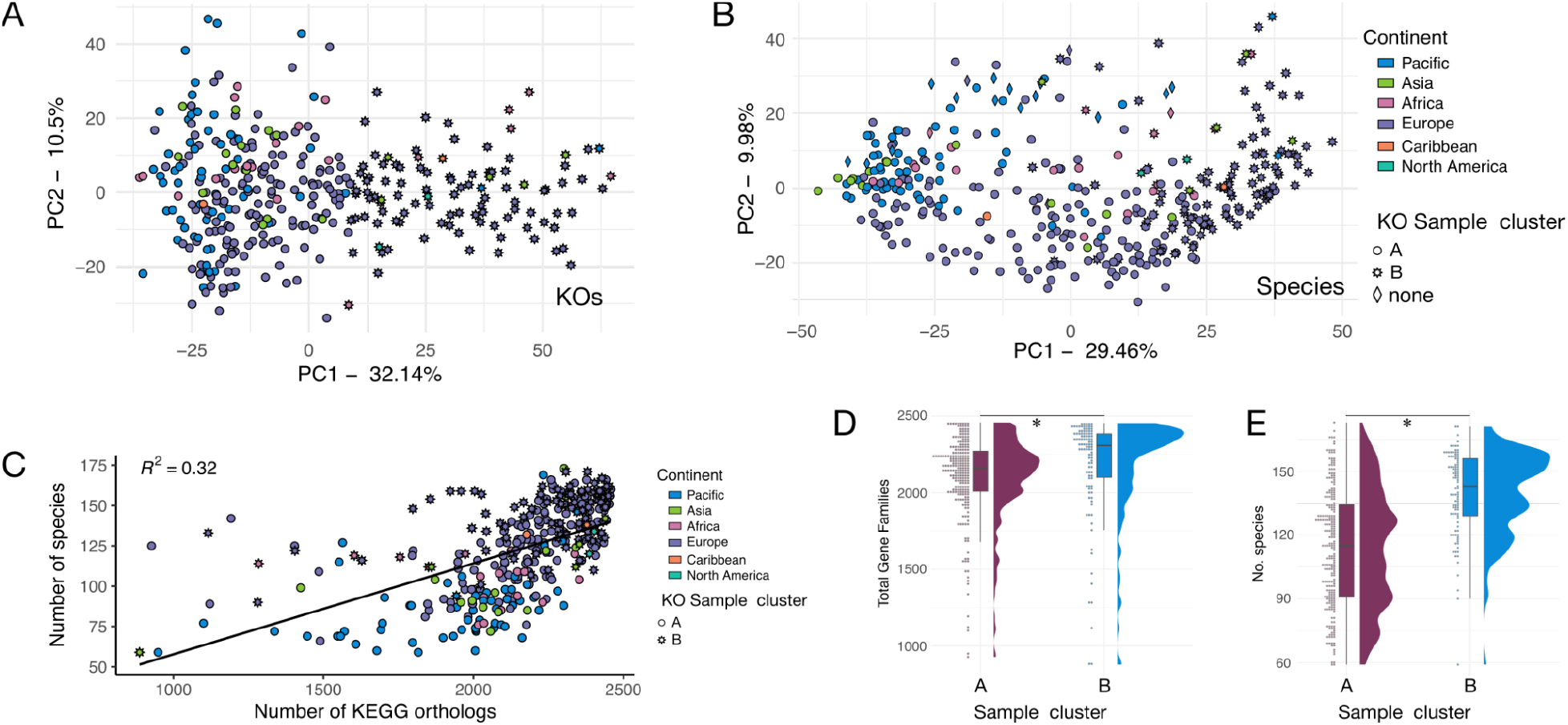
Association between KEGG ortholog counts and species counts in samples. **A.** PCA based on KEGG ortholog abundance, colored by continent and shaped by cluster from hierarchical clustering shown in main Figure 4. **B.** PCA based on species abundance (same as main Figure 3A) colored by continent and shaped by cluster from KEGG ortholog clustering, same as panel A. The diamond samples with no cluster were excluded from KEGG Ortholog analyses. **C.** Correlation between the number of species and the number of KEGG orthologs detected in a sample. **D.** The number of KEGG orthologs in each sample, grouped by the sample cluster from hierarchical clustering. **E.** The number of species in each sample, grouped by the sample cluster from hierarchical clustering. * p < 0.05.

**Supplemental Figure S8.**
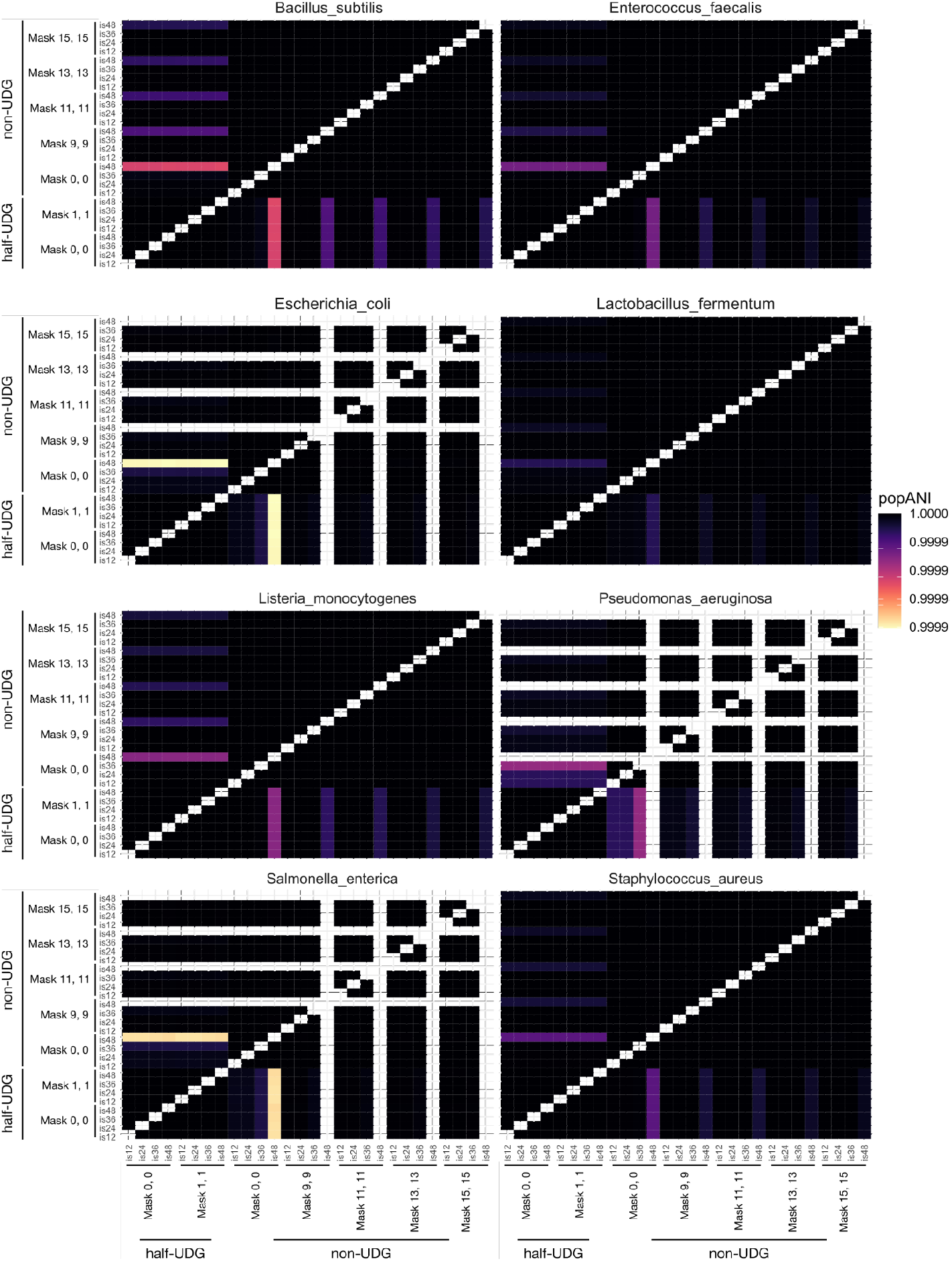
Simulated metagenome parameter testing for inStrain assessment of strain diversity in ancient metagenomes. **A.** PopANI of species in a synthetic community with differing levels of aDNA damage (C-T transitions). Synthetic communities were simulated with high C-T transition rates (non-UDG) or low C-T transition rates (UDG-half). Following mapping against reference genomes, the ends of reads were masked in the mapped read bam file at different lengths along the read to see whether and how much C-T transitions affect popANI estimations. In the sample name, the numbers following “mask” indicate the number of bases from the first and last base on each read were masked, i.e. “non_udg_mask_15_15” indicates the read was simulated with high C-T transition rate and the first and last 15 bases of each mapped read were masked to hide potential damaged sites.

**Supplemental Figure S9.**
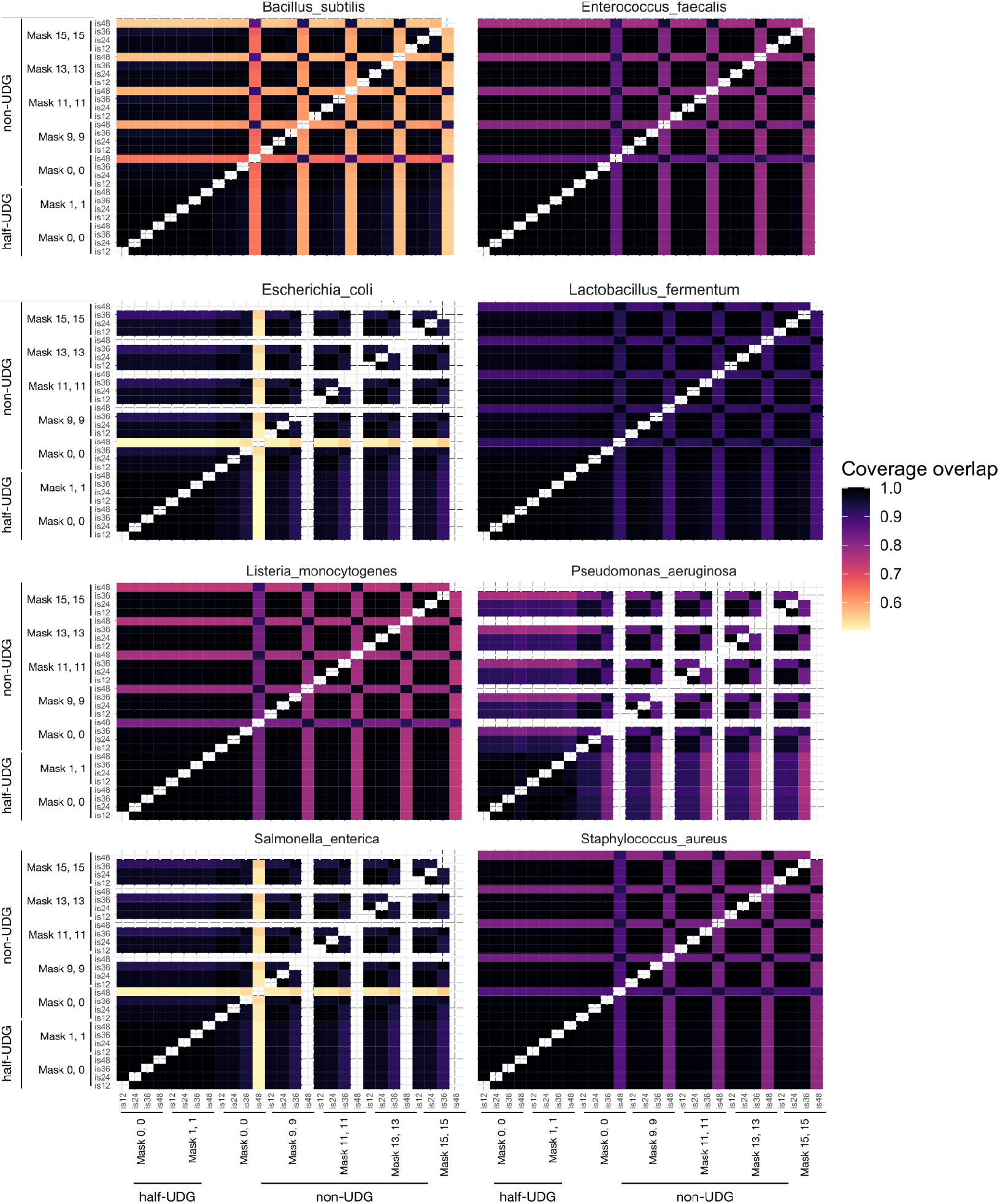
Simulated metagenome parameter testing for inStrain assessment of strain diversity in ancient metagenomes. Percent of genome compared between samples in a synthetic community with differing levels of C-T transitions. Synthetic communities were simulated with high C-T transition rates (non-UDG) or low C-T transition rates (UDG-half). Following mapping against reference genomes, the ends of reads were masked in the mapped read bam file at different lengths along the read to see whether and how much C-T transitions affect popANI estimations. In the sample name, the numbers following “mask” indicate the number of bases from the first and last base on each read were masked, i.e. “non_udg_mask_15_15” indicates the read was simulated with high C-T transition rate and the first and last 15 bases of each mapped read were masked to hide potential damaged sites.

**Supplemental Figure S10.**
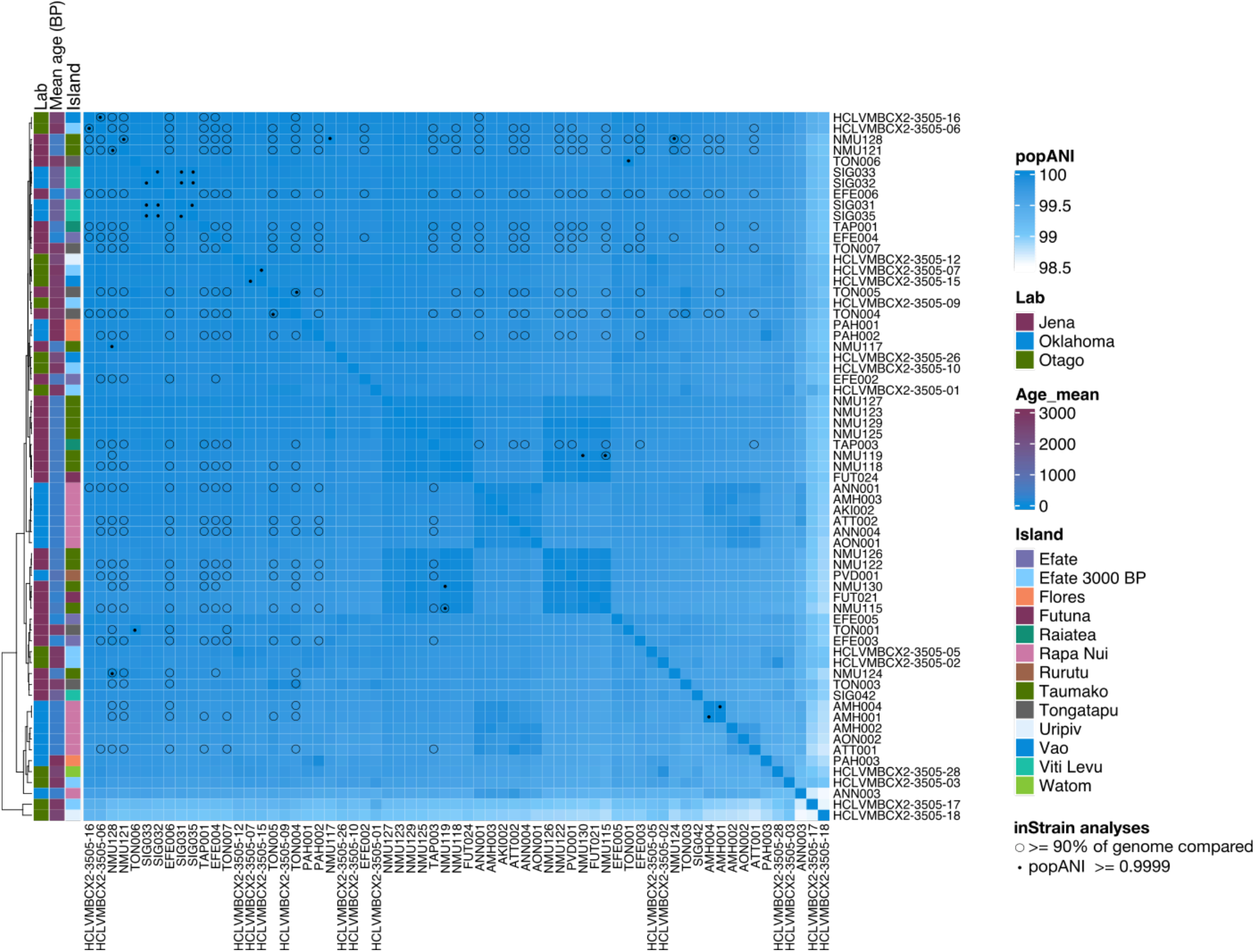
Heat map showing the inStrain popANI for *Anaerolineaceae* bacterium oral taxon 439 between Pacific samples. Black dots indicate samples that have a popANI ≥ 0.9999 (range: 0.9999083 - 0.9999970), and open black circles indicate samples where ≥ 90% of the genome could be compared. Only samples AMH001 and AMH004 have a popANI > 0.99999.

**Supplemental Figure S11.**
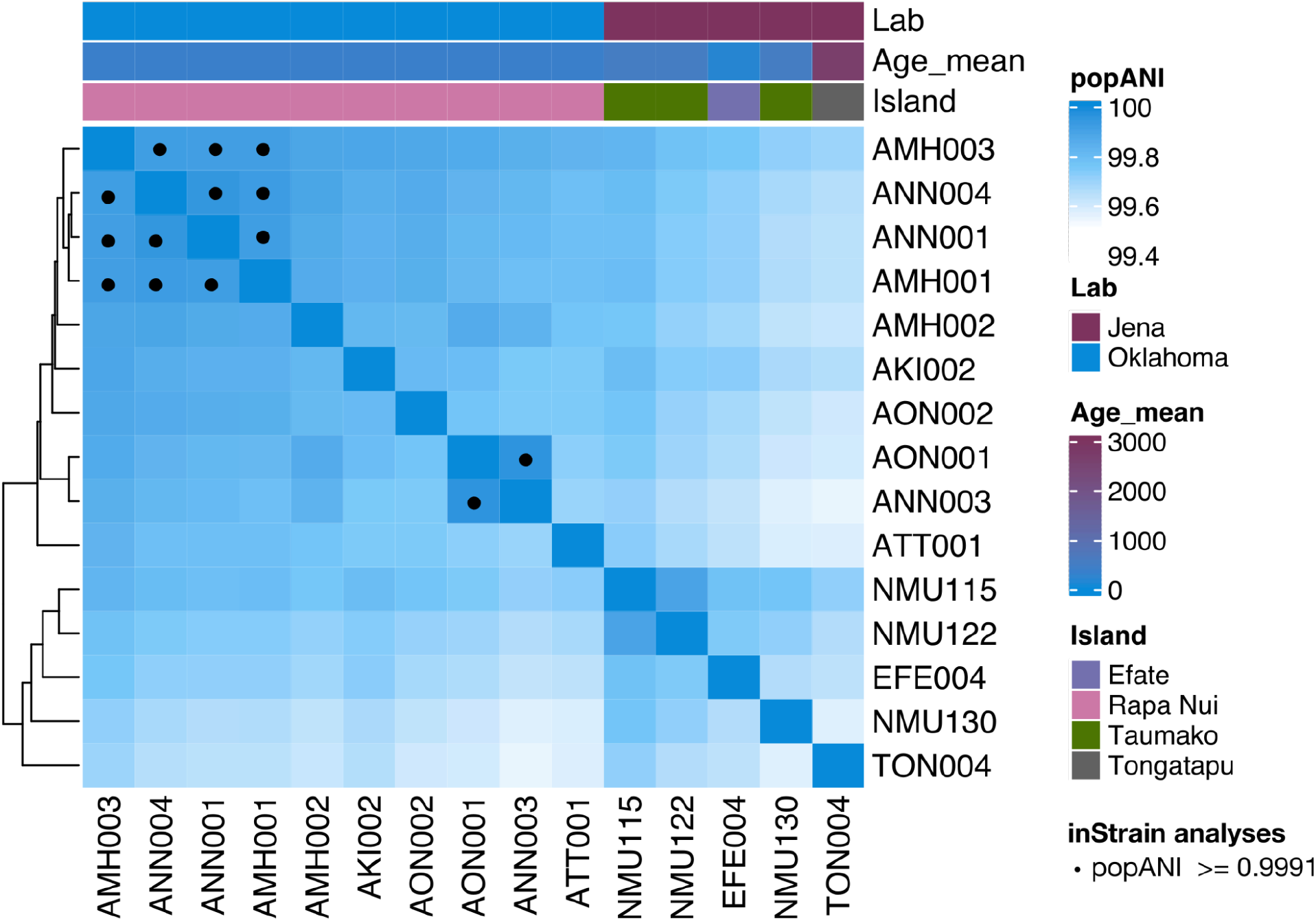
Heat map showing the inStrain popANI for *Tannerella forsythia* between Pacific samples. Samples are hierarchically clustered. The percent of genome compared between each sample was at most 82%, suggesting that low coverage of this species reduces the ability to distinguish strains. Black dots indicate a popANI of > 0.9991 (range 0.9991 - 0.99955).

**Supplemental Figure S12.**
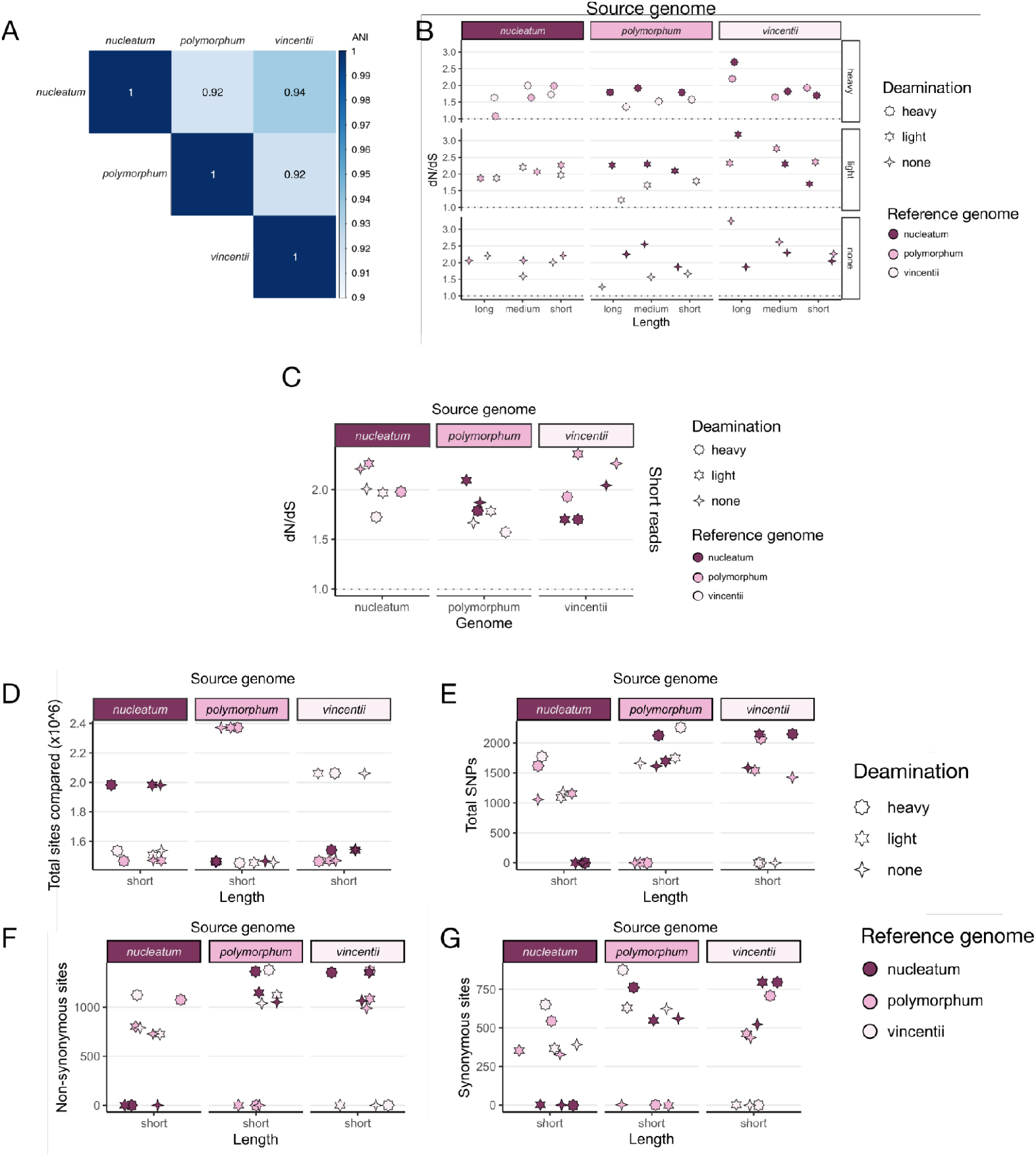
*Fusobacterium nucleatum* subspecies as a test-case for dN/dS values for mapping against an incorrect but closely-related species. Three subspecies of *Fusobacterium nucleatum* (*nucleatum*, *polymorphum*, and *vincentii*) were reduced to short read sequencing data of three lengths, long, medium, or short (150bp, 75bp, or 30bp, respectively), and ancient DNA damage C-T transitions were added in a high or low amount, or left off (none, essentially modern DNA). A. ANI values of each subspecieis genome compared to the other two. B. dN/dS values calculated with polymut after mapping the short read data against each of the three subspecies genomes. C. dN/dS of short read data of each genome mapped against the other genomes. D. Total sites compared for each genome compare to the other genomes. E. Total SNP’s identified by polymut with default parameters. F. Total non-synonymous SNPs for each genome mapped against the other genomes. G. Total synonymous SNPs for each genome mapped against the other genomes.

**Supplemental Figure S13.**
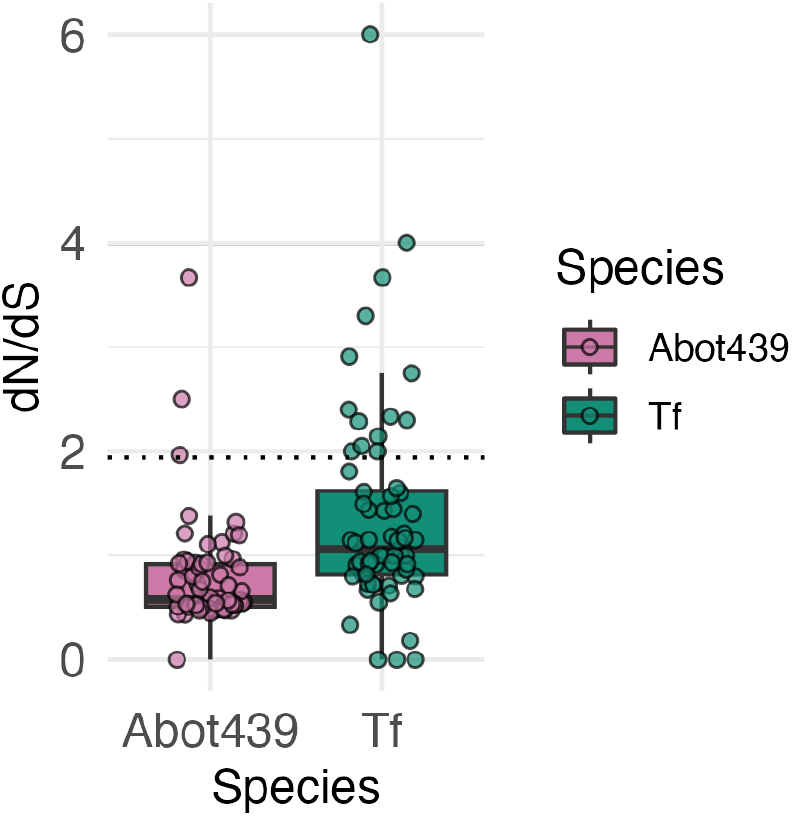
The ratio of non-synonymous SNPs to synonymous SNPs (dN/dS) for all samples mapped against the *Anaerolineaceae* bacterium oral taxon 439 or *Tannerella forsythia* genome. The dotted line at 1.94 indicates the dN/dS of mapping a species to a closely-related reference genome (see Supplemental Figure S12). Abot439 - *Anaerolineaceae* bacterium oral taxon 439; Tf - *Tannerella forsythia*.

**Supplemental Figure S14.**
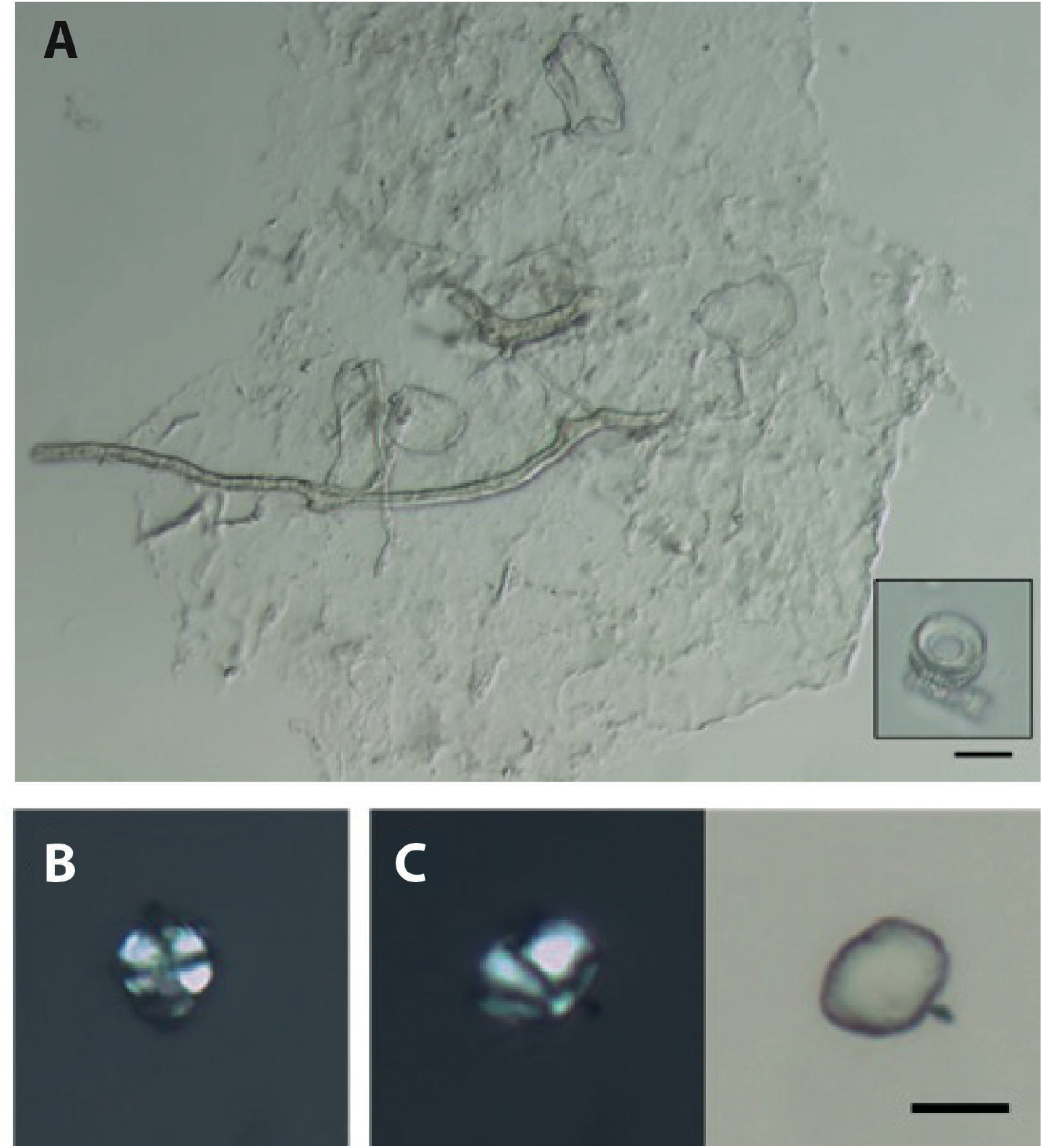
Microfossils observed in dental calculus from Efate. (**A**) Fungal spores and hyphae, with inset showing two diatoms; 10 µm scale bar applies to both images. Starch granules from (**B**) EFE003.B (cross-polarized light) and (**C**) EFE006.B (left in cross-polarized light, right in transmitted light); scale bar is 10 µm and applies to both images.

**Supplemental Figure S15.**
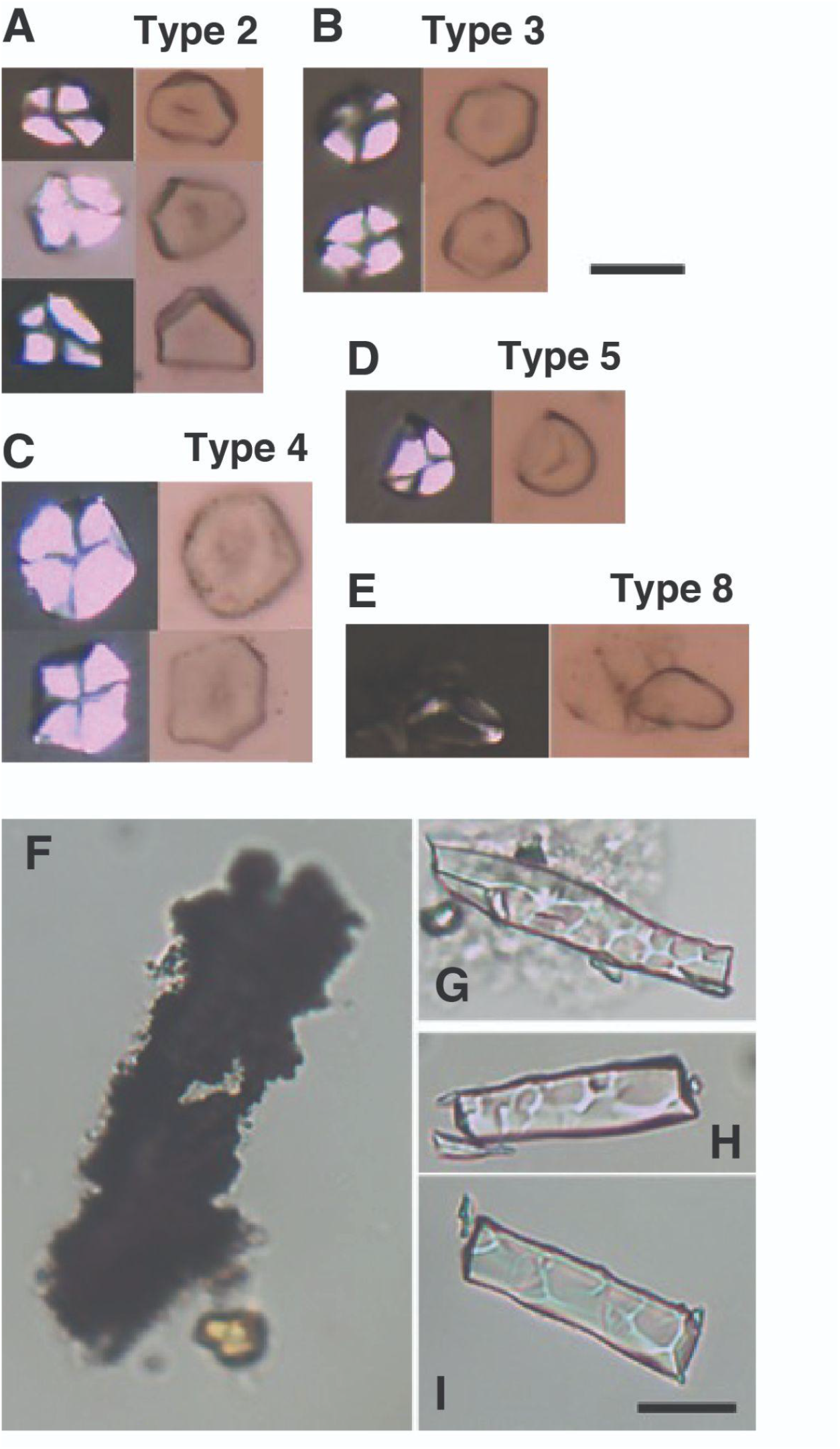
Microfossils observed in dental calculus from Futuna. (**A-E**) Starch granules from sample FUT018.B (left in cross-polarized light, right in transmitted light); scale bar is 10 µm and applies to all images. (**F**) Microcharcoal and (**G-I**) phytoliths from sample FUT021.B; scale bar is 20 µm and apply to all images.

**Supplemental Figure S16.**
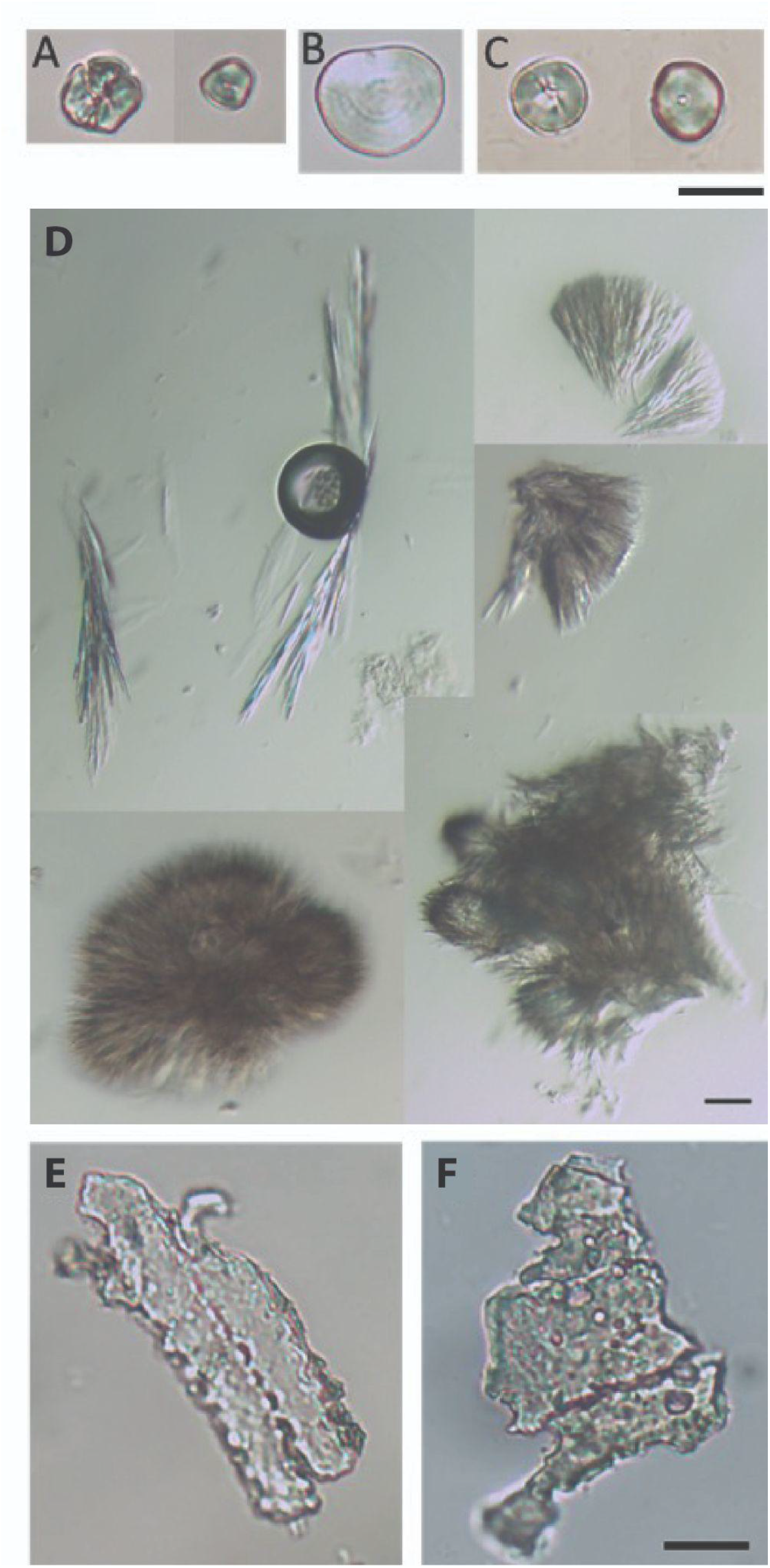
Microfossils observed in dental calculus from Taumako. Starch granules from (**A**) NMU119.A, (**B**) NMU122.A and (**C**) NMU127.A; scale bar is 20 µm and applies to all samples. (**D**) Unknown microparticles from sample NMU116.A; scale bar is 20 µm and applies to all images. (**E**) Dentate elongate phytoliths from NMU122.A, and (**F**) damaged phytoliths from NMU123.A; scale bar is 20 µm and applies to both images.

**Supplemental Figure S17.**
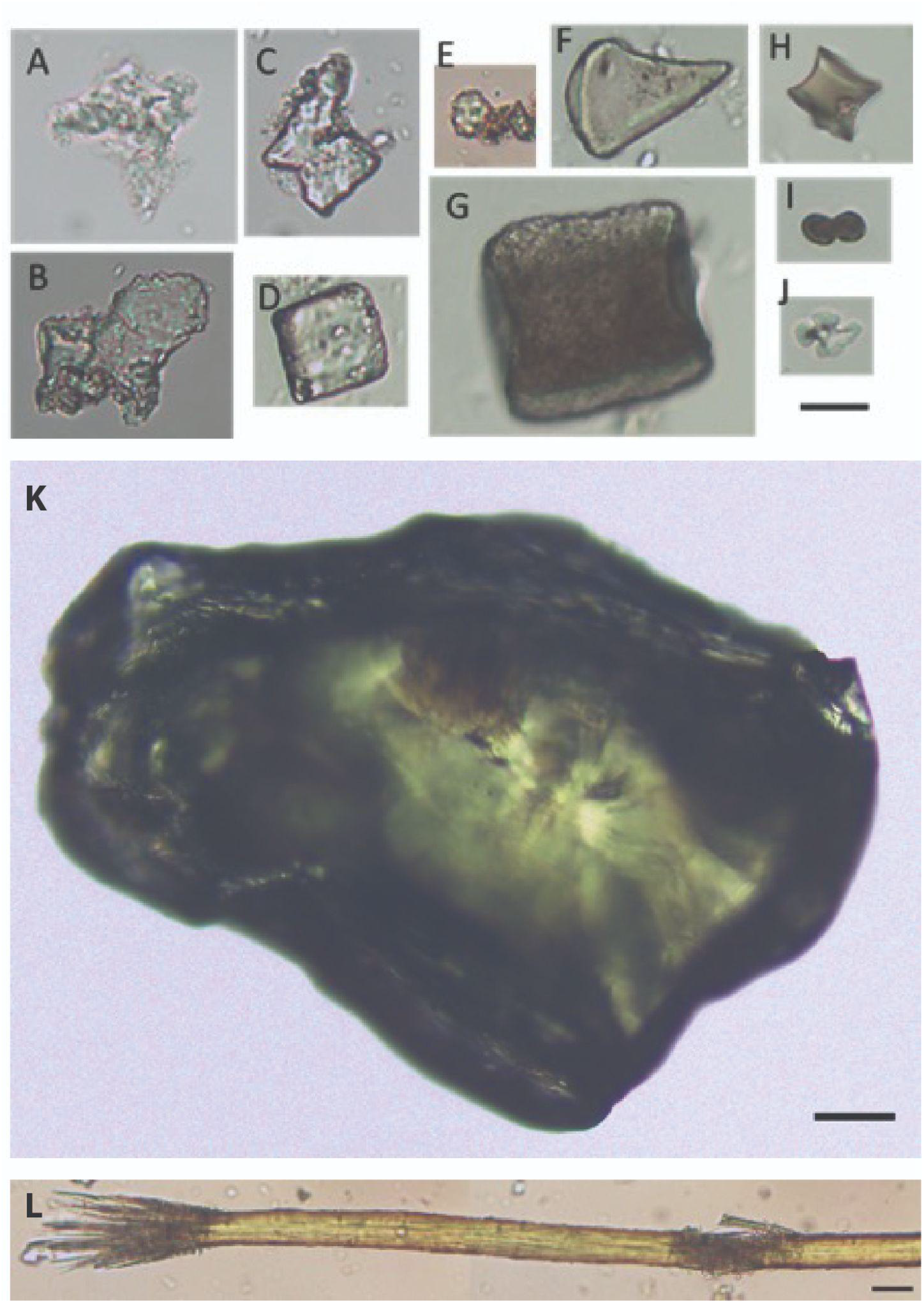
Microfossils observed in dental calculus from Fiji. Phytoliths from SIG040.A (**A, B**), SIG042.A (**C**), SIG031.A (**D**), SIG045.A (**E**), SIG032.A (**F, G**), SIG036.A (**H, I, J**). **A** and **C** look similar to double peaked glume phytoliths; **B** is an amoeboid echinate phytolith of unknown origin; **D** and **G** are blocky type phytoliths; **E** is a spheroid echinate (palm) phytolith; **F** is an acute bulbus pytolith; **H** is a burnt rondell phytolith; **I** is a burnt bilobate phytolith; **J** is a cross phytolith; scale bar is 10 µm and applies to all images. (**K**) Example of the abundant olive green mineral particles found in the Fiji samples; scale bar is 50 µm. (**L**) Fiber found in sample SIG044.A; scale bar is 50 µm.

**Supplemental Figure S18.**
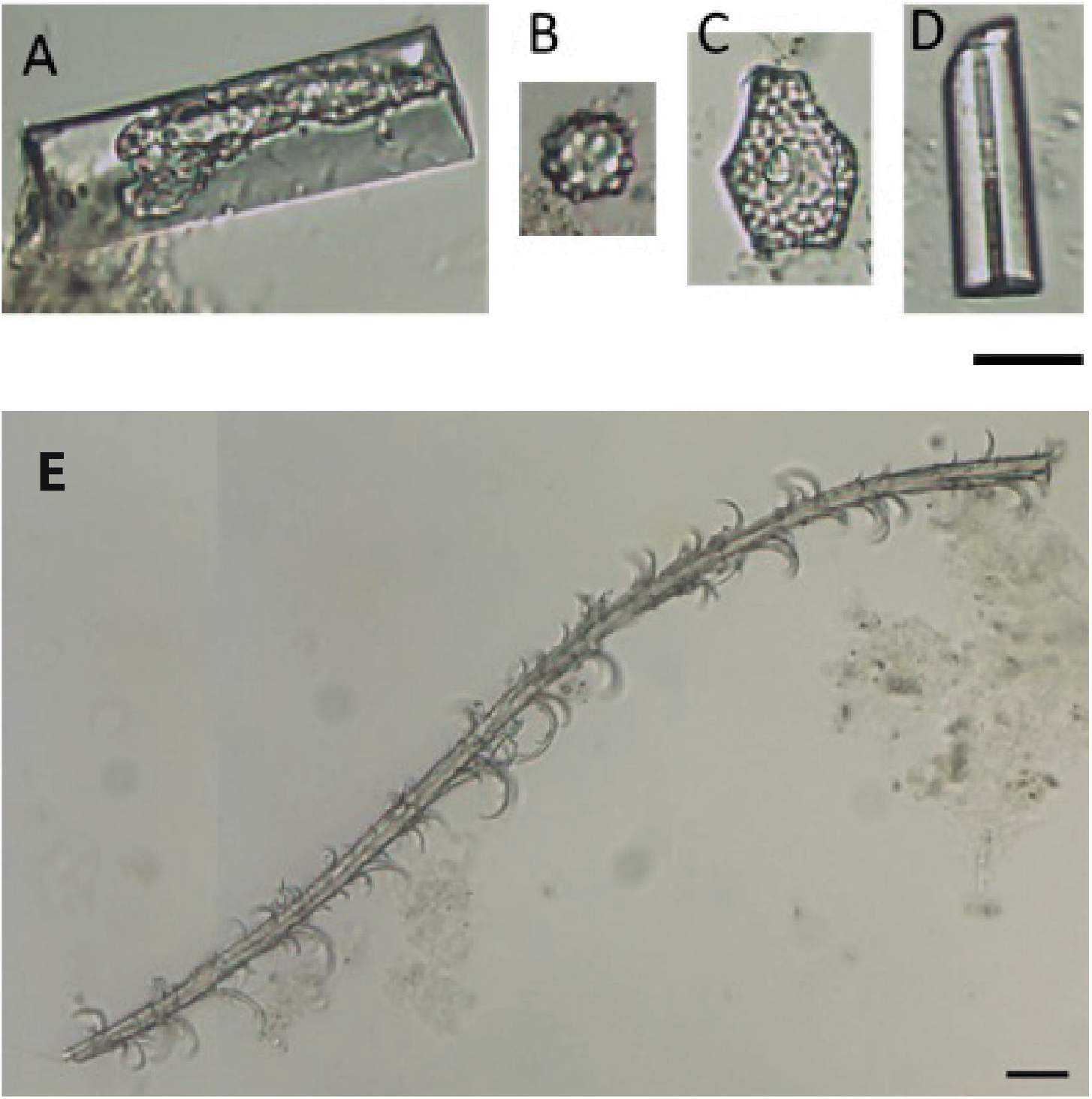
Microfossils observed in dental calculus from Tongatapu. (**A-D**) Examples of phytoliths and sponge spicules recovered from Tongatapu. (**A**) Elongate psilate; (**B**) Spheroid echinate; (**C**) Polygonal scorbulate; (**D**) sponge spicule; scale bar is 20 µm. (**E**) Potential feather barbule from TON001.C; scale bar is 20 microns.

